# Hierarchical structure is employed by humans during visual motion perception

**DOI:** 10.1101/758573

**Authors:** Johannes Bill, Hrag Pailian, Samuel J Gershman, Jan Drugowitsch

**Author notes:** Equal contribution (Samuel J Gershman, Jan Drugowitsch).

## Abstract

In the real world, complex dynamic scenes often arise from the composition of simpler parts. The visual system exploits this structure by hierarchically decomposing dynamic scenes: when we see a person walking on a train or an animal running in a herd, we recognize the individual’s movement as nested within a reference frame that is itself moving. Despite its ubiquity, surprisingly little is understood about the computations underlying hierarchical motion perception. To address this gap, we developed a novel class of stimuli that grant tight control over statistical relations among object velocities in dynamic scenes. We first demonstrate that structured motion stimuli benefit human multiple object tracking performance. Computational analysis revealed that the performance gain is best explained by human participants making use of motion relations during tracking. A second experiment, using a motion prediction task, reinforced this conclusion and provided fine-grained information about how the visual system flexibly exploits motion structure.

---

The visual scenes our brains perceive in everyday life are filled with complex dynamics. Information hitting the retina changes not only with every motion in the scene, but also with every head movement and saccade. Noise, occlusions, and ambiguities further make visual information inherently unreliable. In order to maintain stable, coherent percepts in the face of complex and unreliable inputs, our brains exploit the spatially and temporally structured nature of the environment [1].

Motion structure refers to statistical relations of velocities. One form of structure common in natural scenes is motion grouping: when a rigid object is set in motion, all of its visual features move coherently. This structure allows us to infer the existence of objects based on the coherent motion of features—the Gestalt grouping cue known as *common fate* [2]. Grouping based on common fate has been shown to influence our ability to track objects [3–6], to search displays [7], and to store information in short-term memory [8].

However, the strict definition of common fate is too brittle to accommodate natural scenes in which visual features do not move together rigidly and yet are still grouped together. In some cases this is because we perceive objects as deforming non-rigidly. In other cases, we perceive objects that are hierarchically structured [9–11]: the parts of an object move rigidly *relative* to a reference frame (the object), which itself could be a rigidly moving part of another object, and so on. For example, we perceive the motion of hands relative to the motion of arms, and the motion of arms relative to the motion of the torso (**Fig. 1A,B**). The entire body may be moving relative to another reference frame (a train or escalator). The perception of hierarchically organized motion suggests a powerful “divide-and-conquer” strategy for parsing complex dynamic scenes.

**Figure 1.**
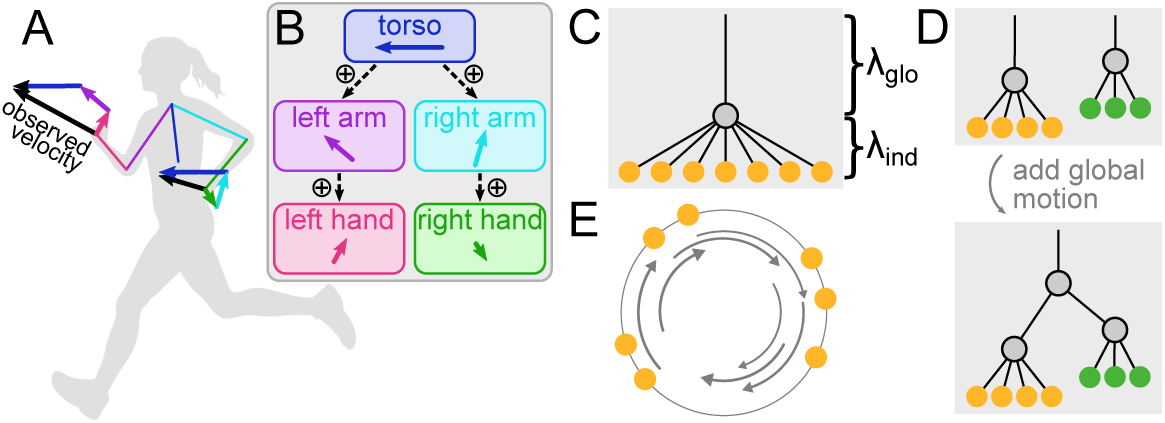
Modular representation of hierarchical motion structure. (*A*) Observed velocity components of a running human. The hands inherit motion from the arms, which inherit motion from the torso. (*B*) Corresponding nested hierarchy of motion relations. Observed velocity is the sum of local motion components. (*C*) Motion graph describing global motion with a strong (strength *λ*_glo_) shared motion source (gray node) and weaker (strength *λ*_ind_) individual motion sources (orange nodes). Here, the global motion source is not directly observed (i.e., latent) but introduces correlations in the motion of observable objects (orange). (*D*) Two motion clusters (top) can be embedded into a deep hierarchy by adding another latent motion source at the tree’s root (bottom). (*E*) Illustration of a stimulus of stochastically rotating dots with global motion structure. These are the class of stimuli used in the experiments.

Recent work has formalized the representation and discovery of hierarchical motion structures [10]. However, we still know relatively little about if and how the visual system exploits this structure for visual tasks like object tracking. We address this question in two experiments, one testing multiple object tracking (MOT, [12]) and one testing multiple object trajectory prediction. A key innovation of these experiments is the use of tightly controlled stimuli that isolate the signatures of hierarchical structure from other forms of motion structure. We show that a Bayesian observer model, when equipped with the appropriate structural representation, can provide insight into the mental computations underlying human behavior across a variety of motion structures. In the MOT task, we find that improvement of tracking performance for structured stimuli cannot be explained without exploiting structure knowledge during inference, indicating that humans make use of motion structure knowledge when perceiving dynamic scenes. In the prediction task, we uncover the employed motion structure knowledge of our participants, finding that humans can flexibly recruit different motion relations for scene parsing, including deep motion hierarchies which nest groups of objects within another moving reference frame.

## Results

The study was pre-registered [13] prior to data collection. All pre-registered analyses are presented in the main text; any additional analyses are labeled as such.

### Representation of motion structure

In this section, we introduce a representation of hierarchical motion structure. To motivate this representation, consider a flock of birds. The velocities of individual animals in the flock are naturally described by the sum of a global motion component, which is shared across all birds, and an individual component for each animal relative to the flock’s velocity. Only the aggregate motion is observed; the underlying motion components are latent variables that help us organize our perception.

To formalize the idea of shared, latent motion components, we assume all velocities in a visual scene to be driven by motion sources (nodes in the graph in **Fig. 1C**) that either represent observable objects (filled nodes) or are latent (unfilled nodes). A motion source inherits velocity from a parent source when connected by an edge, thus supporting tree-like hierarchies. Motion sources can be shared by multiple objects, and the total velocity of an object is the sum of all inherited motions. In addition to the graph connectivity, each motion source has a motion strength *λ* that determines the source’s contribution strength to the speed of dependent observable objects. The simple “global motion” motif in **Fig. 1C** might, for example, to a first order describe the motion structure underlying a flock of birds, where larger motion strengths are illustrated by larger vertical distances (curly braces) between the motion sources.

The separation of motion composition (graph connectivity) and motion strength (vertical node location) gives rise to a flexible, modular representation of motion structure, covering many real-world scenes such as independent motion, clustered motion or deep motion hierarchies. The graph in **Fig. 1D** (top), for example, describes a faster orange and a slower green cluster of otherwise independently moving composed objects (e.g., two non-interacting flocks of birds). A further global motion component, such as an observer moving his head, would introduce another motion source at the root of the tree (see **Fig. 1D** bottom). This illustrates how our motion representation can be used in a modular fashion to describe deep nested motion hierarchies. In fact, any tree-like motion structure built in this way can be represented by a *motion structure matrix* ***L***, accommodating both composition and strengths (see *Material and Methods*), such that we will often refer to specific motion structures by their associated matrices ***L***.

This modular representation enabled us to generate motion-structured visual stimuli, which isolated the computational role of motion structure in dynamic visual scenes, and were thus particularly suited for psychophysics experiments. Specifically, we strove to design stimuli in which all object properties and statistical relations among objects were dominated by *structure in velocities*, while keeping other factors like individual object velocities or spatial structure uninformative. To do so, we generated the velocities of observable objects by random draws from a continuous time stochastic process (namely a multivariate Ornstein-Uhlenbeck process [14]; see *Material and Methods* for details) that yielded smooth random trajectories with the desired statistical motion relations imposed by a chosen motion structure ***L***. In this process, motion sources play the role of random forces which accelerate or decelerate any dependent object. For example, the latent global source in **Fig. 1C** would induce correlations among the velocities of all observed objects. The resulting trajectories are mathematically tractable and feature real-world properties, such as inertia and friction, without making any assumptions about specific trajectory realizations. To further remove any persistent spatial structure among the objects’ locations, stimuli were positioned on a circle (illustrated in **Fig. 1E**; see *SI Appendix* for example video). This makes the locations of objects asymptotically independent. The mathematical tractability of the trajectory-generating process was crucial for precise stimulus control. Knowing the exact joint probability distribution across object velocities allowed us to vary velocity correlations induced by motion structure while keeping the motion velocity statistics of individual objects unchanged, thus making structure in velocities the dominant feature in the presented scenes.

### Motion structure improves visual tracking performance

We first asked whether motion structure knowledge benefits humans in a multiple object tracking (MOT) task [12]. Evidence exists that structure among objects, such as grouping [15], symmetry [6] or global translation [4], impacts tracking performance, but hierarchical structure has not been investigated before.

In the MOT task, *K* = 7 dots rotated about a circle with 3 dots being initially marked as targets. After a few seconds, all dots changed to identical appearance while dot motion continued for another 6 s. After that the dot motion stopped, and participants had to re-identify the initially marked targets. We tested 20 participants on four different blocked motion conditions of 30 trials each, with motion graphs shown in **Fig. 2A** (targets marked by *; see *SI Appendix* for example trial videos). Independent (IND) motion, the standard MOT task, served as a baseline. In the global (GLO) condition, dot motion is composed of a dominating shared stochastic motion source, as well as small individual per-dot motion sources to break emergent spatial patterns. In counter-rotating (CNT) motion, the latent motion source affects dot velocities in opposite directions: similar to gears, the force accelerating the dots *k* = 1, 2, 3 in one direction, accelerates dots *k* = 4, 5, 6, 7 (three of which are targets) in the reverse direction. The last motion condition is a counter-rotating deep hierarchy (CDH) in which two 3-dot groups are driven by a counter-rotating motion source that shares a global motion source with the 7th dot. This deep nested structure generates distinguishable velocity patterns that cannot be approximated by any shallow (max. one latent source layer) motion structure. In the CDH condition, we tested two different target sets (CDH_1_ and CDH_2_) to probe set-specific effects in tracking performance. We measured participant performance as the average number of correctly identified targets within each condition. Since the aim of the experiment was to test how humans can exploit motion structure knowledge rather than learn it, at the beginning of each motion block, participants were explicitly presented with a diagram and three demonstration trials of the motion structure that would be presented in the upcoming block. Overall average stimulus speed was titrated on a per-participant basis to approximately reach a performance of 2.15 (midpoint between chance level (3 × 3/7) and perfect (3)) correctly identified dots in the IND condition. This per-participant speed level was subsequently maintained for the rest of the experiment. During the following data collection, the marginal motion statistics of individual dots were then held constant for each participant across all conditions ***L*** and dots *k* such that conditions only differed in their dot velocity correlations, as determined by ***L***. A separate IND condition block (see left-most bar in **Fig. 2B**) that was not used to adjust the stimulus speed confirmed the validity of the adjustment and marks the reference for performance changes in motion-structured stimulus conditions. The performance on the four conditions with motion structure (GLO, CNT, CDH_1_ and CDH_2_) are shown in **Fig. 2B** (left) next to the IND reference performance. The introduction of structured motion significantly impacted dot tracking performance, (*p* ≈ 1.03 × 10^−14^, Greenhouse-Geisser corrected repeated measures ANOVA) resulting in a significant performance boost in all motion conditions (one-sided paired t-tests, *p* ≈ 1.2 × 10^−8^, 2.4 × 10^−9^, 7.9 × 10^−3^, 2.6 × 10^−4^, respectively; see *SI Appendix*, Fig. S1 for additional pairwise comparison of all conditions). In conclusion, motion structure clearly improved human tracking performance.

**Figure 2.**
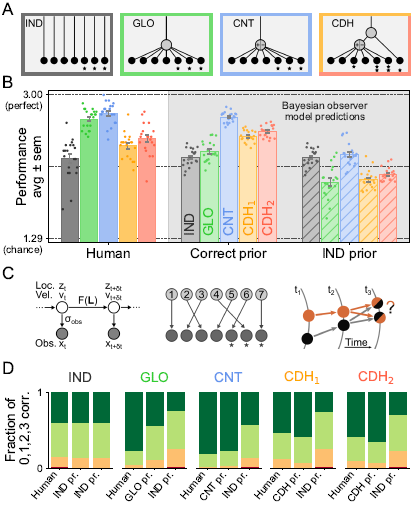
Use of motion structure knowledge during multiple object tracking. (*A*) Tested motion conditions included independent (IND), global (GLO), counter-rotating (CNT), and counter-rotating deep hierarchical (CDH) motion. *’s mark targets. Two different target sets were tested for CDH motion. (*B*) Average performance (number of correctly identified targets) on different motion conditions by human participants, the Bayesian observer model using the correct motion structure, and the Bayesian observer disregarding motion relations (IND prior). Using motion structure during inference is required to explain human performance gains on motion-structured stimuli. (*C*) The Bayesian observer model consists of a Kalman filter with motion structure prior ***L*** (left) and a mental assignment of dot identities (center). Perceptual and neural noise can lead to ambiguous assignments and, ultimately, errors in the reported target set (right). (*D*) Fraction of trials with 0, 1, 2, and 3 dots correct (red, orange, light- and dark-green respectively) for human participants, the observer model with the correct motion prior, and the model with an IND prior.

### Use of structure drives MOT performance gains

The observed boost in tracking performance for structured stimuli could simply be a byproduct of dot velocity correlations, that is, an intrinsic stimulus property, rather than the result of using motion structure knowledge by the observer. For instance, the paths of dots with positively correlated velocities might cross less frequently during a trial, making their confusion less likely [16]. To distinguish the contribution of employed motion structure knowledge from stimulus-intrinsic factors, we extended a Bayesian MOT observer model by Vul and colleagues [17] to incorporate motion structure priors ***L***. The deliberately simple model (illustrated in **Fig. 2C**) includes only the core components required to perform the MOT task, making it a minimalistic, normative model of motion-structured MOT (see *Material and Methods* for details). Similar to Vul et al. [17], visible dot locations ***x***_*t*_ in individual video frames are subject to perceptual and neural noise *σ*_obs_ while being integrated with mental estimates of location ***z***_*t*_ and velocity ***v***_*t*_ via a multi-dimensional Kalman filter (**Fig. 2C** left). To capture structured motion, we extended this Kalman filter to incorporate the motion structure matrix ***L*** as a Bayesian prior distribution of how dot velocities are expected to evolve over time. A global (GLO) motion prior, for example, would favor positively correlated dot trajectories. The Kalman filter alone, however, assumes that every observed dot location is labeled by the dot’s identity, making dot confusions, and therefore imperfect dot tracking, impossible. To produce such confusion, we needed to additionally model the mental assignment of dot identities to the visually identically looking dots on the screen (**Fig. 2C** center, cf. [17]), which is a manifestation of the correspondence problem [18, 19]. Errors in this mental assignment can lead to imperfect re-identification of the target dots at the end of an MOT trial, and arise when dots come close [20] or even cross: uncertainty in internal location estimates ***z***_*t*_ can render multiple mental assignments possible, as exemplified in **Fig. 2C** (right). Since the Bayesian model is derived solely from computational considerations for solving MOT and disregards any psychophysical constraints, e.g., velocity-dependent observation noise [21], we refer to it in what follows as the *computational observer model*.

We presented the computational observer model with the exact same trials that had been shown to human participants. The single free parameter of the model, *σ*_obs_, was adjusted to match average human performance in the baseline IND motion condition, and was subsequently fixed to predict performance in all other conditions. This resulted in a qualitative match between human performance and the observer model’s predictions (**Fig. 2B** left vs. center). Besides the motion structure dependence, the observer also correctly predicted the increased performance for the non-split target set (CDH_2_) over the split set (CDH_1_). For a discussion of the quantitative mismatch in the GLO condition, see *“Probing components of human object tracking”*, below, and *SI Appendix*.

More importantly, the computational observer allowed us to distinguish whether the performance gain for structured motion stimuli was simply due to stimulus-intrinsic factors: if some motion structures were generally easier to track than others, the computational observer should be able to take advantage of these differences even without explicitly exploiting the structural knowledge in its prior. To test this, we repeated the above procedure with an observer that assumed an IND motion prior, which respects the correct marginal dot velocities, but disregards any knowledge on velocity correlations, even when motion structure was present in the stimuli. As **Fig. 2B** (right vs. left) reveals, this modification caused all motion structure-dependent performance gains to vanish. Hence the observed performance gain of human participants cannot be explained by stimulus-intrinsic factors alone. Instead, humans appear to make use of motion structure knowledge when performing the task.

The finding that the Bayesian observer with correct motion prior explains large parts of human responses, while the motion structure-agnostic observer (IND prior) does not, is further supported by comparing the fraction of MOT trials with 0, 1, 2, or all 3 dots identified correctly (**Fig. 2D**, additional analysis). This visual impression is corroborated by pre-registered statistical measures: while the cosine similarity between per-participant human and correct-prior observer model performance gains (“structured condition minus IND condition”) is highly significant (*p*10^−5^, one-tailed test against H0: positive cosine similarity is a random effect), the one between per-participant human and IND-prior observer model performance gains is not (*p* ≈ 0.99).

### Probing components of human object tracking

The modularity of the Bayesian observer affords insights into the differential role of different psychophysical constraints impacting MOT performance. Specifically, we examined two alternative observer models which add/remove components to/from the computational observer (see **Fig. 3A**; additional analysis, details in *Material and Methods*).

**Figure 3.**
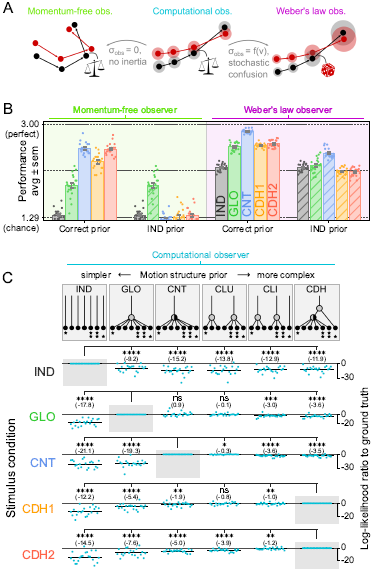
Functional components underlying human MOT. (*A*) Alternative observer models with components added to or removed from the computational observer. The “momentum-free observer” lacks the concept of inertia. The “Weber’s law observer” adds two known psychophysical constraints: velocity-dependent observation noise and stochastic decision noise. (*B*) Average MOT performance on different motion conditions for the alternative observer models when either the correct motion structure or an IND prior are assumed (like Fig. 2B). Employing trajectory extrapolation is computationally indispensable on the circle, as highlighted by the momentum-free observer’s performance dropping to chance in the IND stimulus condition. Adding psychophysical constraints, in contrast, leads to an improved match with human performance. (*C*) Bayesian model comparison of employed motion structure priors based on the exact per-trial choice sets. Shown are log-likelihood ratios of human choice sets under different putative motion priors ***L*** relative to the true structure underlying the stimulus (see main text for description). Negative values indicate that the participant’s behavior was better explained by the correct motion prior. Human participants make use of motion structure knowledge during tracking, presumably employing approximately correctly structured priors. Yet, the limited information provided by the discrete response sets prevented insight into the exact structural features used. One dot per participant, stimulus condition and putative motion prior. Horizontal lines show mean log-likelihood ratios across participants (values in parentheses). *‘s indicate significance of paired t-tests (*p* 0.05, 0.01, 10^−3^ and 10^−4^ respectively).

- The *momentum-free observer* lacks the concept of inertia, i.e., that objects usually change their trajectory smoothly. This model mirrors the perceptual assumptions from established work [10] where motion relations give rise to only correlated location displacements of objects between stimulus frames. We further removed any observation noise, *σ*_obs_ = 0, to isolate the effect of inertia for MOT.
- For the *Weber’s law observer*, in contrast, we augmented the computational observer with two key psychophysical constraints modeled in [17], namely, speed-dependent observation noise [21] and decision noise [22, 23] in dot assignments.

The MOT performance of both models is shown in **Fig. 3B** when either the correct prior or an IND prior are assumed. We find that the momentum-free observer’s performance drops to chance level in the IND condition, confirming that the use of inertia is crucial for performing MOT in the circular task design: whenever two dots come close in one frame, they can only be tracked to the next frame by exploiting the fact that trajectories continue smoothly. This insight complements previous studies of 2D motion tracking [17, 24] for which very close dot proximity was less likely, and hence the use of trajectory extrapolation less critical. In fact, this might have been the reason why previous studies found little evidence of use of inertia in MOT. However, when structure is present, as seen in the CNT and CDH conditions, even a momentum-free observer can benefit from motion structure knowledge for partial performance recovery. Notably, such motion prior-dependent performance gain is not present when tracking GLO stimuli, pointing to a peculiarity of this stimulus condition (see *SI Appendix*, for a computational discussion based on the dot proximity auto-correlation function, Fig. S4). The Weber’s law observer, on the other hand, benefits from structured priors in all conditions and predicts human responses better than the computational observer (cf. also *SI Appendix*, Fig. S2 for trial speed-dependent performance and Fig. S3 for exact choice sets), underscoring the contribution of known psychophysical constraints to human visual tracking.

A second direction supported by the Bayesian observer is a detailed analysis of how well the different models predict the exact set of dots chosen by human participants (note that these are additional analyses, see *SI Appendix*). For probing the putative use of motion prior ***L***, we used simulations to estimate the likelihood of choosing a particular set of dots in each trial of the experiment for observer models with different motion priors. We then asked how well these simulated likelihoods match the participants’ choices. Results of this analysis are shown in **Fig. 3C** for the computational observer, and in *SI Appendix*, Fig. S3 for the momentum-free and Weber’s law observers. This analysis confirms the above result that some motion structure knowledge is employed by human observers, in that an IND prior explains human responses significantly worse than the respective correct motion priors. Besides the structures used in the stimulus conditions, the comparison includes two additional priors: clustered motion (CLU) and clusters-plus-independent (CLI) motion. These priors approximate the motion relations of the CDH condition without considering its nested nature. This analysis suggests that human participants typically employ the correct or close-to-correct motion priors, even for nested structures—yet, the effect size is often too small to make decisive statements.

In conclusion, the MOT task revealed that humans make use of motion structure knowledge during visual tasks. However, the limited information provided by the participants’ responses (three chosen dots per trial) did not allow us to identify which exact motion features were employed.

### Revealing the structure of human motion priors

We therefore developed a second experiment, multiple object prediction (pre-registered), that grants a more fine-scaled insight into the cognitive machinery underlying motion perception.

Multiple object prediction uses partially observed dynamic visual scenes to test human perception in face of uncertainty. As before, in each trial, seven dots rotated stochastically about a circle (see **Fig. 4A**). After 5 s, however, two target dots (the larger green and red dots) became invisible while the motion of the other dots remained visible. After another 1.5 s the scene froze, at which point participants had to predict the location of both target dots with a computer mouse. We used highly volatile motion-structured stimuli with quickly changing dot velocities, such that the use of motion structure information was indispensable for making predictions. We further colored the dots according to their role in the motion structure to prevent dot confusion. In contrast to previous single-dot prediction tasks [11], the rationale for simultaneously predicting two dot locations is that the covariance pattern of errors convey additional information about the motion structure assumptions of our participants. For example, if global motion was assumed, we would expect the participants to jointly over- or underestimate the red and green targets’ final locations.

**Figure 4.**
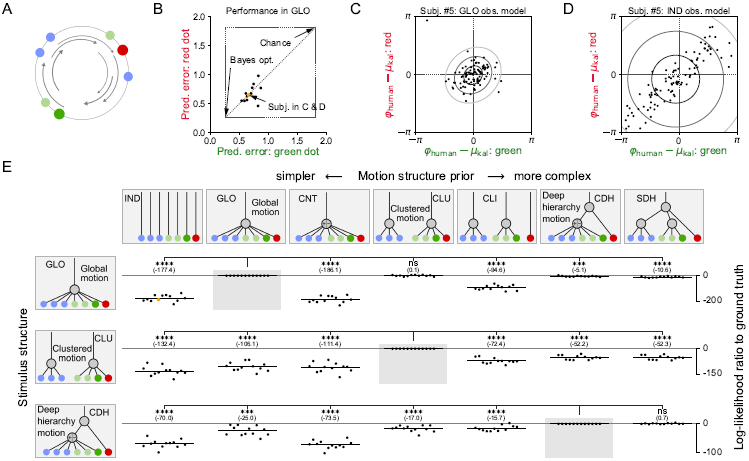
Revealing human motion priors in a multiple object prediction task. (*A*) Illustration of the stimuli. The highlighted green and red dot disappeared after 5 s. Participants had to predict their location at the end of the trial. Dots were color-coded to indicate their role in the motion structure. Here, a global (GLO) stimulus condition is illustrated. (*B*) Mean squared prediction error of the green and red dot for all participants in the GLO condition. Due to task-inherent uncertainty, even an ideal observer exhibits non-zero prediction errors. Humans do not reach the accuracy of the ideal observer, but perform better than chance. (*C*) Human responses (dots) relative to the predictions of an ideal observer with correct GLO prior, in all 100 trials for the participant highlighted in B and E. The fitted observer model (ellipses) predicts human responses well (ellipses indicate 1, 2, 3 standard deviations). (*D*) Same as C, but for an observer model assuming an IND motion prior. Neither the predicted locations nor the covariance in human prediction errors is captured by the model. (*E*) Left: tested motion conditions included global (GLO), clustered (CLU) and counter-rotating deep hierarchical (CDH) motion. Top: putative motion priors tested for explaining human responses via a Bayesian observer model, ordered by their complexity (see main text for description). Each cell shows per-participant log-likelihood model fit ratio for a particular motion prior, compared to the correct prior underlying the stimulus (indicated by gray background). Negative values indicate that the participant’s behavior was better explained by the correct motion prior. Humans flexibly employed correctly structured motion priors. One dot per participant, stimulus condition and putative motion prior (comparison between C and D highlighted in orange). Horizontal lines show mean log-likelihood ratios across participants (values in parentheses). *’s indicate significance of paired t-tests (*p* 0.05, 0.01, 10^−3^ and 10^−4^ respectively).

We tested 12 participants on 3 motion conditions each: global motion (GLO), clustered motion (CLU) and counter-rotating deep hierarchical motion (CDH) (motion graphs in **Fig. 4E**; example videos in *SI Appendix*), with 100 trials per condition. As before, participants were briefly trained on all motion structures and were informed about the specific structure underlying each trial. Participants reported the task to be challenging, but performed reasonably well (see **Fig. 4B** for the GLO condition, and *SI Appendix*, Fig. S8 for CLU and CDH; additional analysis).

In order to identify the structure of the motion prior employed by each participant in each stimulus condition, we formulated a Bayesian observer model of human responses in the prediction task. This model was the same as our MOT model, but without the dot confusion component, and with location observations assumed to be practically noise-free (*σ*_obs_ = 0, except for the invisible dots for which no further observations were provided). After each trial, motion-structured Kalman filters with different putative motion structure priors ***L*** predicted the statistically optimal, most likely location of the target dots based on the motion of the other dots, and the uncertainty in this prediction. Building on [22], we linked this prediction to the locations reported by the participants by assuming additional response variability that scaled with prediction uncertainty (correlated across the two target dots), motor noise (uncorrelated across dots), and the possibility to confuse the red and green dots when reporting their locations (see *Material and Methods* for details). The first two variability sources were modeled as Gaussians, owing to the Gaussianity of observed human response error distributions (see *SI Appendix*, Fig. S6; additional analysis), whereas the third variability source was implemented by a small swapping probability. Overall, this led to three model parameters that we fit by maximum-likelihood separately for each participant, condition, and putative motion structure prior. Applied to simulated behavior with the same trials, we found that this procedure was able to correctly recover the motion structure underlying simulated responses (*SI Appendix*, Fig. S7).

To identify the most likely motion structure prior employed by the participants, we compared across 7 structurally different putative motion priors how well the model can explain the participants’ trial-by-trial predictions. For the GLO motion condition, the model predictions are shown in **Fig. 4C,D** when assuming a GLO prior (C) or an IND prior (D) for all 100 trials of one representative participant (see *SI Appendix*, Fig. S9 for all participants and stimulus conditions). The visually apparent better match of the GLO over the IND prior is statistically quantified by the log-likelihood ratio of the human responses under the two motion priors (see **Fig. 4E**, highlighted data point in the top left matrix cell). The resulting log-likelihood ratios for all motion conditions and priors are shown in **Fig. 4E**, with putative motion priors sorted from simpler to more complex structures (pre-registered; see *SI Appendix*, Fig. S10 for additional comparison across a larger range of motion structures, yielding similar results). Note that all compared models have the same number of free parameters. For the global (GLO) motion condition (first row of **Fig. 4E**), human responses are best explained either by the correct global motion prior or by more complex priors which would be similarly suited for solving the task (e.g., because they contain a global motion source). Motion priors without a global motion source, such as IND, CNT, and CLI, resulted in significantly worse model fits. For the clustered (CLU) motion condition (second row), all alternative model priors led to worse model fits. Together, these results establish confidence in the applicability of the Bayesian observer model.

Our most complex motion condition, the counter-rotating deep hierarchy (CDH; **Fig. 4E** bottom row), was designed to require inference over both latent motion sources for correct prediction. In other words, none of the tested shallow (non-nested) motion priors can mimic the error covariance pattern expected under a CDH prior. Global (GLO) motion would miss the counter-rotating component of the blue and green groups. Counter-rotating (CNT) motion would miss the global motion component. A clusters-plus-independent (CLI) motion prior (see **Fig. 4E** top) could approximately predict the green dot’s location, but lacks the global component for predicting the red dot. The only structures that could approximate a CDH covariance pattern are other deep nested hierarchies, such as a standard deep hierarchy (SDH) that employs a third latent motion source. The result of Bayesian model comparison (bottom row in **Fig. 4E**) clearly favors deep hierarchical motion priors for explaining human predictions. Overall, our results strongly suggest that human observers are able to use motion structure, including hierarchically nested motion relations, when perceiving dynamic scenes.

### Systematic and stochastic errors underlie human sub-optimality

Even though participants employed the qualitatively correct motion structure in the prediction task, they featured additional, suboptimal variability when compared to a statistically optimal observer (cf. **Fig. 4B**). This variability could arise from consistent deterministic bias, indicating a systematic mis-integration of the stimuli, or stochastic fluctuations, indicating potentially noisy computations. To determine the contribution of each alternative to overall variability, we repeated each trial twice within each condition, supporting a bias-variance-decomposition [23] of this variability (**Fig. 5A**). Specifically, if all of the variability was due to deterministic biases, participants should feature the exact same, systematic errors, Δ^(1)^ = Δ^(2)^, in both trials. If, in contrast, all variability was purely stochastic, the observed errors should be uncorrelated across paired trials. This idea allowed us to quantify the ratio between bias and variance by one noise factor *f*_noise_ per participant and motion condition, with *f*_noise_ = 0 and *f*_noise_ = 1 denoting purely bias-driven and purely noise-driven observers, respectively (see *Material and Methods* for details). We evaluated the noise factor separately for the green and red target dot, with the results shown in **Fig. 5B**. Note that these results are additional analyses. We found that human sub-optimality is a combination of systematic and stochastic errors. The noise factors of both dots were on average equally strong (black **×**). The positive correlation between red and green dot noise factors (*ρ* = 0.51, Pearson correlation) suggests a participant-dependent level of “noisiness”. This impression is further supported by consistently higher or lower noise factors across participants (colors & filling in **Fig. 5B**; *p* 10^−4^, one-way Welch’s ANOVA).

**Figure 5.**
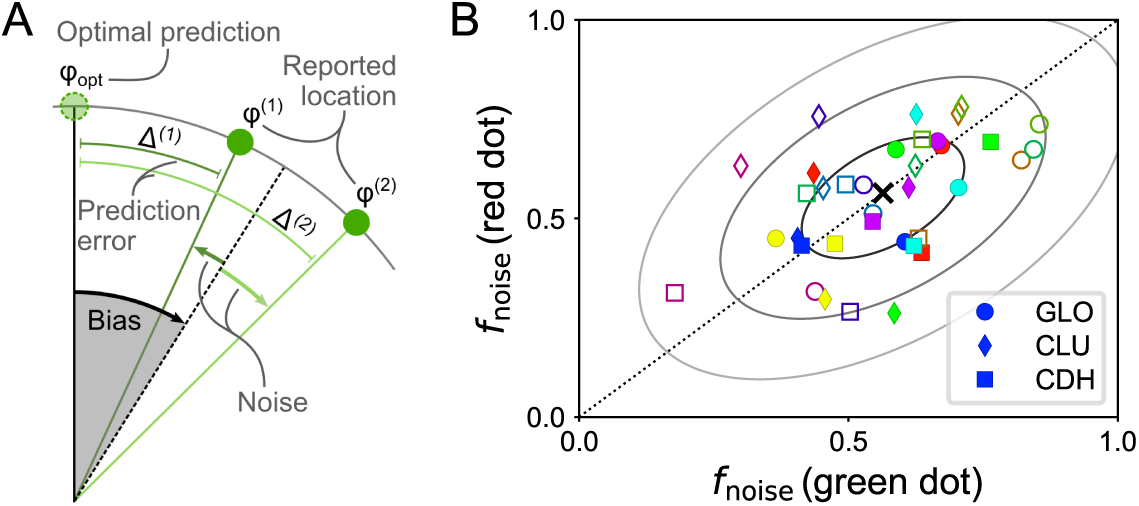
Bias-variance decomposition for the prediction task. (*A*) Human prediction errors are assumed to be the sum of a systematic (bias) and a stochastic (noise) component. The relative contribution of both components can be estimated from repetitions of the same trial, leading to responses *φ*^(1)^ and *φ*^(2)^, and associated errors Δ^(1)^ and Δ^(2)^. (*B*) Noise factors for the green and the red dot, one marker per participant (color & filling) and motion condition (shape). A combination of systematic and stochastic errors underlie human sub-optimality. Black cross and ellipsoids indicate mean and iso-density curves (1-3 SDs) of a bivariate Gaussian fitted to all noise factors.

## Discussion

We have shown that humans make use of motion structure knowledge during demanding perceptual tasks, and that they can flexibly employ structured motion priors for the task at hand. Beyond shallow motion motifs, such as global motion or clustered motion, we were able to show that humans can even use deep nested motion hierarchies in the prediction task. Key to revealing the covert motion priors of human participants was the design of analytically tractable stimuli that capitalize on motion relations. Analytical tractability facilitated the development of normative Bayesian observer models, our mathematical scalpel for dissecting the use of motion priors by the participants from stimulus-intrinsic contributions to visual motion perception.

Understanding motion perception as Bayesian inference has a rich scientific history [10, 11, 17, 25–29]. Bayesian models of low-level perception [25–27] were extended to explain human multiple object tracking of independent dot movement [17, 29]. Our work extends this line of research to structured motion, revealing the use of structured motion priors during tracking. The modularity of the observer model further enabled us to study the role of specific computational components, such as employing only sub-trees of a motion structure, as well as the role of specific perceptual components, such as noise at different stages or exploiting the concept of inertia during tracking. We did not consider further features underlying human performance, such as attention effects [15, 29, 30], tracking of subsets of dots, or not consistently employing all features of a motion structure, as these are more challenging to operationalize within our framework, and were not central to this study. A systematic deviation between human MOT performance and model predictions in the GLO condition finally points to additional temporal integration on intermediate time scales (∼1 s; cf. *SI Appendix* Fig. S4), going beyond established explanations based on object distance [20, 31].

Our aim to resolve the fine structure of human motion priors has thus led us to the development of a novel probabilistic multiple object prediction task that augments single object prediction [11] with velocity covariances within a tractable protocol. Including posterior covariance matrices in the observer model greatly enhanced the log-likelihood ratios in the prediction task beyond the resolution achievable in single object prediction (see *SI Appendix*, Fig. S11), an effect potentially related to self-consistency biases [32]. While we had originally developed our modular matrix representation ***L*** of hierarchical structure as an extension to the motion trees in [10], the proposed decomposition into graph connectivity and motion strengths bears some resemblance to the semantic structure interpretation of singular value decomposition applied in [33] to categorical data and to Pythagorean tree embeddings used in [34] for describing language structure. Flexible structure representations are expected to foster cognitive science research on how humans infer the motion structure of a visual scene—potentially recruiting from a set of motion features—and how such motion features for modular combination could be learned in the first place.

Biologically, the importance of motion for visual scene perception is reflected in the tuning of cells in primate visual areas. Neurons in area MT are frequently tuned to the speed and direction of velocity within their receptive field [35], while downstream areas, like MSTd, encode progressively richer motion primitives such as selective tuning to expansion, rotation and spiraling [36, 37]. This points to a feature repertoire that is tailored to behaviorally relevant stimuli such as radial expansion (the visual pattern on the retina of a forward-moving observer) or rotation (when tilting your head to the left/right). Little, however, is known about how such motion primitives are further recruited in neural circuits for high-level, hierarchical motion processing. Our motion structure representation is compatible with existing neural implementations of Bayesian sensory integration, such as the neural Bayesian filtering model of Beck et al. [38]. In combination with physiology-based models of motion integration [39, 40], this could bring forth normative neural models of structured motion perception in higher visual areas, and guide experiments on the neural code along the visual motion pathway.

## Material and Methods

### Motion structure matrix representation

We describe the motion of *K* visible objects that are driven by *M* motion sources. Usually, *M K* since, besides shared latent sources, each object can feature individual motion. We represent how motion source *m* affects object *k* via a composition matrix of motion motifs ***C*** ∈ ℝ^*K*×*M*^. If *m* drives *k*, we set *C*_*km*_ = 1; if *m* does not affect *k*, we set *C*_*km*_ = 0; for counter-rotating motion, we set *C*_*km*_ = +1 and −1 for opposing directions. Thus, each column of ***C*** encodes a motion motif (e.g., all 1s for global motion), which is tied to motion source *m*. Each motion source has an associated real-valued motion strength *λ*_*m*_ *≥* 0, and we define the motion structure matrix ***L*** as the product of composition and strengths, ***L*** = ***C* Λ** with **Λ** = diag(*λ*_1_, .., *λ*_*M*_), i.e., strength *λ*_*m*_ scales the *m*-th motion motif.

### Motion-structured stimuli

The stochastic dynamics for location ***z***_*t*_ ∈ [0, 2*π*)^*K*^ and velocity ***v***_*t*_ ∈ ℝ^*K*^ are given by

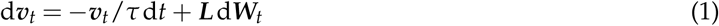

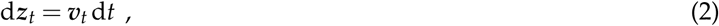

with friction time constant *τ* ∈ ℝ^+^ and *M* independent Wiener processes in vector ***W***_*t*_. The values of *τ* and **Λ** used in each experiment are provided in *SI Appendix*. For simulations, we employed Euler–Maruyama integration to advance the dynamics, and we map ***z***_*t*_ *↦*(***z***_*t*_ mod 2*π*) after each step to keep locations on the circle. The stationary distributions under above dynamics are (see [14]):

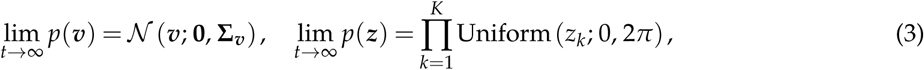

where 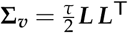, and the expression for *p*(***z***) assumes that each object has a non-vanishing independent motion component. Knowing the stationary velocity covariance **Σ**_***v***_ in closed form gives convenient control over the stimulus. For example, the marginal velocity distribution of the *k*-th object is 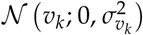 with 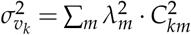·which is simply the sum of all parent motion sources’ squared strengths. In all experiments, we initialize the stimuli from a sample of their stationary distribution. Then, velocities follow a multivariate Gaussian at any time, and locations become asymptotically independent.

### Experiment details: MOT task

Twenty Harvard University undergraduates (Mean age=20.6 years, SD=2.01; 14 females) with normal or corrected-to-normal vision participated in the experiment for course credit. All participants provided informed consent at the beginning of the experiment. The experiment lasted approximately 2 hours and was comprised of a thresholding phase and a testing phase. Across both phases, participants performed a multiple object tracking (MOT) task, in which they were presented with a 2000 ms static display wherein 7 colored dots encircled by white outlines appeared around a larger black ring. All dots proceeded to move around the ring for 5000 ms, following predetermined trajectories. Stimulus motion trajectories were pre-computed using custom Python code and initial values for ***v*** and ***z*** were drawn from the stimulus motion structure’s stationary distributions [3]. When pre-computing the trials, we asserted that the final dot locations were non-overlapping to prevent selection ambiguity (trials with overlap were regenerated until they met this criterion). Red square outlines subsequently appeared for 3000 ms around 3 of the moving dots, marking them as to-be-tracked targets. The target cues and disc colors then faded to black during the following 700 ms, after which the white outlines continued to move for 6000 ms until coming to a stop. A random number (1-7) was subsequently superimposed on each dot, prompting participants to report the identities of the perceived targets by making the appropriate keyboard presses. Feedback was provided after every trial. The MOT tasks implemented across both phases were nearly identical, with the following exceptions. Whereas movement trajectories in the thresholding phase followed independent (IND) motion, those implemented in the test phase varied across blocks (Latin-square counterbalanced) based on each motion structure condition (IND, GLO, CNT, CDH1, CDH2). The thresholding phase was used to titrate motion speed by adjusting a per-participant speed factor *f*_speed_ (see eq. [2] in *SI Appendix*), such that all participants would achieve a common baseline on IND trials. To this end, participants performed 30 IND trials at an initial speed of *f*_speed_ = 2.0. Whether performance at this initial speed fell above or below the targeted 2.15 performance threshold, participants completed a subsequent 30 trials of the same task at a faster or slower speed (speed factors varied by 0.25 increments). This staircase thresholding procedure was repeated until each participant’s average target identification accuracy approximated 2.15 items per trial (Mrepetitions=2.40, SD=0.68). Once thresholding was complete, participants performed 150 trials of the MOT task at their determined speed. Prior to the onset of each motion structure block, participants were presented with three example trials inherent to the forthcoming block of displays, and were provided with a motion graph that explicitly laid out the motion structure based on the colors of the moving items. Example trials and details of experiment conduction are provided in *SI Appendix*. No data was excluded from the analysis. The study was approved by the Harvard Institutional Review Board (IRB00000109).

### Bayesian observer model: Kalman filtering (both experiments)

We assume independent Gaussian observation noise on the locations of each video frame, 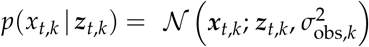, for all dots *k* = 1..7. Together with the latent stochastic linear dynamics of [1] + [2], Kalman filters are the statistically ideal observer models for the stimulus. The circular support of ***z***, however, renders the normal distribution underlying Kalman filtering into an approximation. To maintain a close approximation to the correct posterior, we ensured that variances of the location posterior *p*(***z***_*t*_ |***x***_1:*t*_) remained small at all time. For all observable dots, location estimates are much smaller than *π* anyway; for the unobserved dots in the prediction task, we designed the stimuli such that the correct posterior’s standard deviation never exceeded 40°. Thus, most of the probability mass stayed within the (unwrapped) circle at any time, and errors introduced by the Gaussianity assumption are small. The corresponding matrices of the motion-structured Kalman filter are provided in *SI Appendix*.

### Bayesian observer models: MOT task

Above Kalman filter was used for tracking, assuming either the correct motion prior ***L***^⋆^ to underlie the stimulus, or an alternative putative motion prior ***L*** with identical marginal velocity distributions 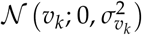. Following [17], different possible mental dot assignments *γ* are scored at each frame by comparing the likelihoods of the current observation ***x***_*t*_ under the possible assignments. As in [17], this poses a numerically intractable inference process (per frame, there are *K*! possible assignments; the total number of assignments per trial thus grows as *K*!^#frames^). We use discrete particle variational inference [41] for approximate inference, with a single particle and the set of all pairwise dot permutations as differential proposals per frame. For the computational and momentum-free observers, after each frame (at time *t*), the highest scoring candidate assignment *γ* is maintained with the trajectory likelihood *p*(***x***_1:*t*_ |*γ*) as score function; for the Weber’s law observer, the assignment is sampled from *p*(*γ*) ∝ *p*(***x***_1:*t*_ |*γ*)^4^. The assignment *γ* at the end of the trial determined which three input dots were chosen as targets. Note that the observer models are not statistically “ideal” observers since they rely on approximate inference for dot assignments and subsume the effect of noise at all computational stages within a single fitted parameter *σ*_obs_. In the numerical evaluation, the Kalman filter was provided with noisy observations, 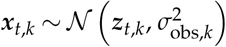 mod 2*π*, for the computational and Weber’s law observers; the momentum-free observer received noise-free observations. The size of *σ*_obs,*k*_ was determined via computer simulations to yield on average the target performance (2.15) in the IND condition. For the computational observer, *σ*_obs,*k*_ = 0.05 for all dots *k* = 1..7; for the Weber’s law observer, *σ*_obs,*k*_ = *σ*_0_ (1 + 5 |*v*_*t,k*_|) and *σ*_0_ ∈ [0.008, 0.016] depending on the speed level *f*_speed_ of the participant. Each trial presented to a human participant was then simulated 25 times (with changing noise instantiations), and simulated performance was averaged over all repetitions to reduce the variance of the observer model’s predictions (shrinking the error bars in the center and right panel of **Fig. 2B** and in **Fig. 3B**). The generative model of the momentum-free observer as well as details on the log-likelihood estimation for the additional analysis in **Fig. 3C**, which is based on the simulated repetitions trials, are provided in *SI Appendix*.

### Experiment details: prediction task

Twelve adult individuals (Mean age=31 years, SD=9 years; 10 males), participated in exchange for financial compensation ($10 per hour plus a performance-dependent bonus). All participants reported normal or corrected-to-normal vision, and provided informed consent at the beginning of the study. On average, the experiment lasted approximately 80 minutes. Each participant completed 100 trials per motion structure (GLO, CLU, CDH), presented in blocks (50 unique trials per block, each presented twice in randomized order). Block orders were balanced across participants. Participants were informed about the stimulus condition and performed a variable number of training trials prior to each block until they decided to start the experiment block. Dots were color coded as shown in **Fig. 4E**. Trials were composed as follows. Initial values for ***v*** and ***z*** were drawn from the stimulus motion structure’s stationary distributions [3]. After a 1000 ms still period, all dots started moving stochastically for 5000 ms according to [1] + [2]. During the end of the 5000 ms period, the red and green target dots faded out, and only the remaining five dots were visible for another 1500 ms period after which the scene froze. The green and red dot’s locations had to be predicted by directing a green / red mouse cursor to the predicted location on the circle. After each trial, the true dot locations were revealed and participants received points (0 – 20) based on the accuracy of their prediction. The points only served as task engagement and for payment, and played no role for the analysis. Example trials and details of experiment conduction are provided in *SI Appendix*. No data was excluded from the analysis. The experiment was approved by the Harvard Institutional Review Board (IRB15-2048).

### Bayesian observer model: prediction task

Kalman filters with different candidate motion priors ***L*** were presented with the same trials that had been shown to human participants. For the observer model, we assume correct dot assignment (*γ* = ***I***) and set *σ*_obs_ = 0 for observed dots since dot confusion and observation noise–induced errors are expected to play a negligible role in the prediction task. While invisible, we set *σ*_obs_ *→* ∞ for the green and red dots. For candidate prior ***L***, human responses 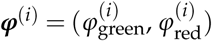 in trial *i* = 1, .., 100 are then modeled as

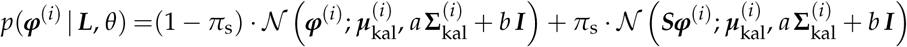

where 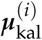 and 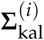 are the (2-dim.) mean and (2 × 2) covariance matrix of the Kalman filter with motion prior ***L*** for the green and red dot at the end of the trial. The matrix 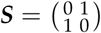 models dot swapping in the response with probability *π*_s_, and *a, b* ∈ ℝ^+^ scale structured inference noise and unstructured motor noise respectively. Eq. **??** describes a *stochastic posterior* observer model (the winning model identified in the systematic comparison of [22]) with lapses and motor noise. The three free parameters *θ* = (*π*_s_, *a, b*) were fitted via maximum likelihood for each candidate prior ***L***, stimulus condition ***L***^⋆^ and participant. In **Fig. 4C,D**, only the non-swapped model component is illustrated. The log-likelihood ratios to the model with correct motion prior,

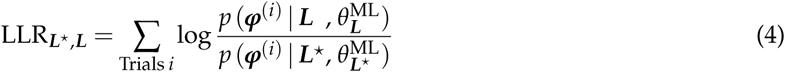

are plotted in **Fig. 4E**. One-tailed paired t-tests identify if the correct prior ***L***^⋆^ explained human responses better than alternative priors ***L***.

### Distinguishing systematic and stochastic errors in the prediction task

We evaluated the noise factor *f*_noise_ separately for the green and red dot per stimulus condition and participant, that is, each noise factor is based on the repetitions *r* = 1, 2 of the unique trials *j* = 1, .., 50 in one stimulus block. We define the average noise variance

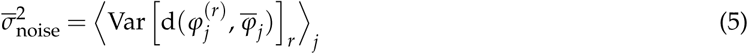

where ⟨·⟩ _*j*_ denotes the average over unique trials, Var[·]_*r*_ is the unbiased estimator of the variance over trial repetitions, d(·, ·) ∈ (−*π, π*] is the (signed) distance on the circle, and 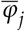 is the circular mean of the two responses 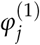 and 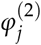. We further define the total prediction variance

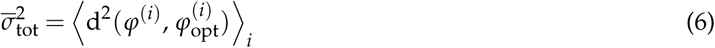

as the mean squared error made compared to the optimal prediction 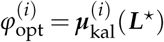. The estimators 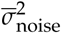 and 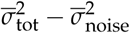 were verified to recover the stochastic and systematic error components of synthetic data (not shown). The noise factor is defined as the ratio 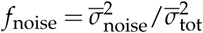. Based on synthetic data, the standard deviation of the noise factors in **Fig. 5B** is estimated to be 0.10.

## Data and code availability

Computer simulations and data analysis were performed with custom Python code which is available on https://github.com/DrugowitschLab/motion-structure-used-in-perception. In this repository we also provide the raw data collected in the experiments.

## Acknowledgments

The authors are very thankful to Rick Born, Till Hartmann, Luke Rast, Ariana Sherdil and Valentin Wyart for helpful feedback and discussions, as well as to George Alvarez for allowing us to conduct the MOT experiment in his laboratory. This research was supported by a seed grant from the Harvard Brain Initiative, an Alice and Joseph Brooks Fund Fellowship award (JB), a research fellowship from the Alfred P. Sloan Foundation (SJG), and a James S. McDonnell Foundation scholar award (Grant #220020462, JD). Parts of this research were conducted using the O2 High Performance Compute Cluster at Harvard Medical School.

## Appendix

### Details on the stimulus generation and Kalman filters

#### Equations of motion with volatility and speed factors

For transparent stimulus control, it is convenient to write the stochastic stimulus dynamics as

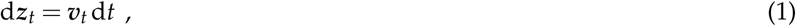

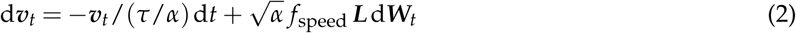

with *speed factor f*_speed_, *volatility factor α* and *M*-dimensional Wiener process ***W***_*t*_. The Wiener process is characterized by increments *W*_*t*+*δt*_ − *W*_*t*_ that are normally distributed with zero mean, *µ* = 0, and variance *σ*^2^ = *δt* (for a discrete-time computer implementation, see below). The speed factor *f*_speed_ scales the absolute value of velocity in the stationary distribution while maintaining the time scale *τ* of typical changes in the stimulus. The volatility factor *α*, in contrast, adjusts the time scale *τ*, while keeping the stationary velocity distribution unchanged, because:

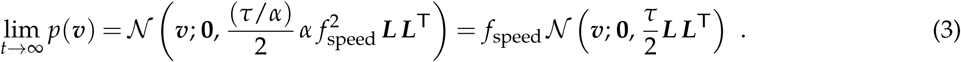

In computer implementations, *f*_speed_ and *α* can be absorbed in *τ* and *λ* via *τ ↦τ*/*α* and 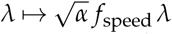. For the MOT task, we adjusted *f*_speed_ in the IND condition in increments of 0.25 until participants performed approximately at the desired target (2.15). After this adjustment, participants covered speed factors between 1.75 and 2.75. All subsequent experiment blocks then used the per-participant speed factor.

#### Parameters of the motion structures used in the experiments

##### MOT task

In the MOT task, we set *τ* = 64s, leading to slowly changing velocities. For *f*_speed_ = 1, the motion strengths in the stimulus conditions were as follows. For independent (IND) motion, *λ*_ind_ = 0.125 for all seven individual sources. For global (GLO) motion and counter-rotating (CNT) motion, *λ*_lat_ = 0.1244 for the latent source and *λ*_ind_ = 0.0125 for the seven individual sources. For counter-rotating deep hierarchical (CDH) motion, *λ*_glo_ = 0.1016 for the global motion source, *λ*_cnt_ = 0.0718 for the counter-rotating sub-source, *λ*_ind_ = 0.0125 for the individual motion sources of the six dots in the clusters, and *λ*_mav_ = 0.0729 for the seventh dot (the Maverick dot). The motion strengths in the CLU and CLI priors for the additional analysis in Fig. 3C match the respective strengths in the GLO condition. Note that motion strengths add up quadratically in the marginal dot velocities due to the factor 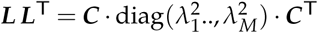. At the example of GLO and IND, we have 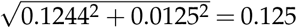 such that the marginal velocities are identical across motion structures. The ***L***-matrices of all conditions are provided in **Fig. 13**.

##### Prediction task

In the prediction task, we set *τ* = 1s, leading to volatile stimuli. The values of *λ* for the various motion sources are provided in **Fig. 10**. The ***L***-matrices of all conditions and observers are provided in **Fig. 14**.

#### Discrete time integration and Kalman filter matrices

For numerical integration, we concatenate (***z***_*t*_, ***v***_*t*_) to a single state vector and advance [1]+[2] of the main paper in small time steps *δt*:

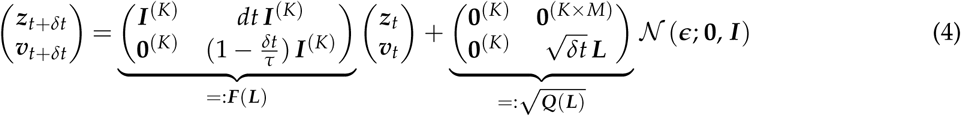

with ***ϵ*** = (*ϵ*_1_, .., *ϵ*_*K*+*M*_) being i.i.d. standard Gaussian noise. The matrices ***F*** and 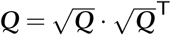 are known as *state transition model* and *process noise covariance* in Kalman filtering.

#### Computer simulations

Stimulus videos where presented at 60 Hz (MOT task) and 50 Hz (prediction task), thereby defining *δt* as the inverse of this value during tracking with Kalman filters. For the generation of stimuli with smooth trajectories, we used an integration time step that was 10-times smaller than the video frame rate for MOT, and an integration time step of 1 ms for the prediction task. Simulations and data analyses were performed with Python 3.6, Matplotlib 3.0, and Scipy 1.2.

#### Log-likelihood estimation for the MOT model comparison

For the model comparisons in Fig. 3C and Fig. 3, we estimated the log-likelihood log *p*(*S*^(*i*)^ = *s*^(*i*)^ |***L***) for the observer model to choose the same set *s*^(*i*)^ as a human participant in trial *i*. For each trial and candidate employed motion structure ***L***, *p*(*S*^(*i*)^ | ***L***) is a categorical distribution with 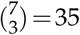 possible values of *S*. The parameters 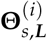 of each categorical distribution were inferred from *R* = 25 repeated computer simulations of the trial. The simulations yielded counts *c*^(*i*)^(*s*, ***L***) of how often *s* was the outcome of the simulation (each repetition had different noise instantiations). To avoid the problem of vanishing probability estimates due to a finite number of repetitions, we put a Dirichlet prior with hyperparameter *α* = 1 on the parameters 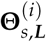. This conjugate prior effectively adds a pseudo-count of 1 to the count *c*^(*i*)^(*s*, ***L***) of every possible set *s*. The likelihood *p*(*S*^(*i*)^ = *s* | ***L***) is then given by the posterior predictive:

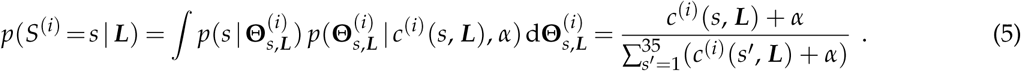

Each point in Fig. 3C and Fig. 3 is the sum over trials of the log-likelihood ratio against the ground truth structure ***L***^⋆^, for one participant:

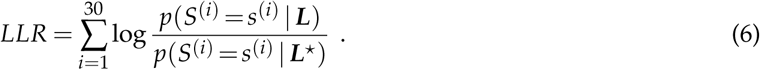

### The alternative momentum-free and Weber’s law observer models

#### Momentum-free observer

The momentum-free observer does not maintain a model of smooth velocity changes. Instead, like in [1], motion relations between dots arise from correlated location displacements. This leads to the following assumed generative dynamics for the momentum-free observer model:

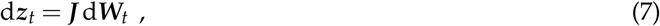

with *K* × *M* structure matrix ***J***.

How does ***J*** have to be chosen to mimic the velocity relations from [1]+[2]? To answer this question, note that, in the stimulus videos, observations ***z***_*t*_ are presented only at a finite frame rate 1/*δt*. We therefore consider the velocity ***v***_*t*_ := (***z***_*t*+*δt*_ − ***z***_*t*_)/*δt*, as it arises from the process [7] when evaluating the difference quotient of subsequent frames. Our goal is to identify ***J*** such that 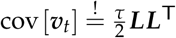, where the r.h.s. is the velocity covariance of the original model. To achieve this, we use that cov[***W***_*t*+*δt*_, ***W***_*t*_] = *t* · ***I*** and that

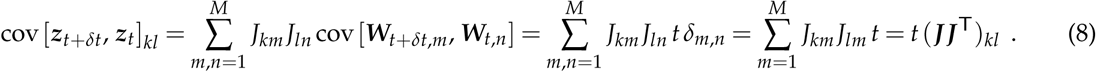

Calculating the covariance cov[***v***_*t*_] is now straight-forward:

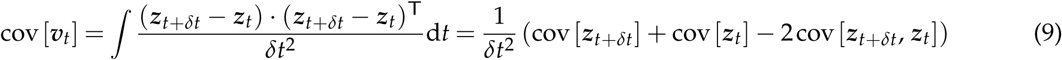

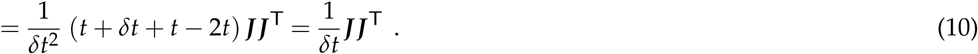

Thus, we can match the motion structure of the original model by setting 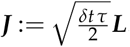. For the discrete time Kalman filter implementation, we obtain (in similarity to [4]) state transition matrix ***F*** = ***I*** and process noise covariance 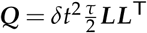.

#### Weber’s law observer

The Weber’s law observer model has been described almost entirely in *Material and Methods* of the main text. Here, we provide only two brief clarifying remarks.

First, the velocity-dependent noise of scale *σ*_obs,*k*_ = *σ*_0_ (1 + 5 |*v*_*t,k*_|) is applied to the observed locations. While velocities are not directly observable in our model, and have to be inferred from noisy locations, the noise (indirectly) also affects the inferred velocities in a velocity-dependent manner. An exact calculation of this indirect effect is tedious—particularly for time-changing velocities. Yet, a naive estimate can be obtained by considering the differential quotient ***v***_*t*_ := (***z***_*t*+*δt*_ − ***z***_*t*_)/*δt* of consecutive frames. When ***z***_*t*_ and ***z***_*t*+*δt*_ are corrupted by independent noise of scale *σ*_obs_(***v***), then ***v***_*t*_ has noise 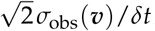, i.e., velocity uncertainty “inherits” the linear ***v***-dependence from the location noise (in accordance with [2]).

Second, we assume that the observer model has knowledge of its location noise during filtering, that is, the size of *σ*_obs,*k*_ is calculated and assigned for each dot. Hereby, the assignment between noise values *σ*_obs,*k*_ and dot identities is done according to the candidate *γ*-permutations.

### Choice of the noise models in the MOT and prediction tasks

The types of noise considered in the observer models are sometimes different for the MOT and the prediction task. In the following, we address why these modeling decisions were made.

#### Observation noise

Observation noise is a prerequisite for dot confusion in the model of MOT. In the prediction task, in contrast, observation noise will only lead to (very small) errors in the inferred locations of the visible dots. The impact of these errors on the inferred locations of the invisible dots is negligible. To keep the model simple, we thus set *σ*_obs_ = 0 in the prediction task observer.

#### Dot confusion

MOT tests dot confusion, and thus this mechanism was included in the analysis. In the prediction task, the risk of dot confusion was greatly reduced due to color coding the dots. The task itself did not rely on dot confusion at all (participants reported the expected location of the invisible dots, which can not be confused with visible dots). Therefore, we did not consider the computationally complex and expensive mechanism of dot confusion in the prediction task observer.

#### Swapping the order of responses

In MOT, swapping the order of keyboard presses does not change the result because only the set of dots was evaluated (not the order of reporting). In the prediction task, the order does matter (which dot is reported as the green and the red one) and is therefore included in the analysis.

#### Decision noise

For MOT, the role of decision noise is not critical for the computational description and, thus, not included in the computational observer. The effect of decision noise is included in the Weber’s law observer model. In the prediction task, responses are continuous, and thus even small decision noise will affect the response. Accordingly, decision noise is included in the analysis.

#### Motor noise

In MOT, participants reported the perceived dot assignment by keyboard presses, for which motor noise should have a negligible effect. In the prediction task, in contrast, responses are continuous and reported by moving the mouse to the desired location. Even small motor noise will affect the response and, thus, motor noise is included in the analysis.

The overarching reason for these choices is to keep the models simple whenever there is good reason to believe that excluding a noise type will not have substantial impact on the result of the analysis.

### Details to the experiments

#### MOT task

The MOT tasks used in both thresholding and test phases were displayed using MatLab Psychophysics Toolbox on a 21.5” iMac monitor (viewable area: 10.54 cm x 18.74 cm, refresh rate: 60 frames/sec). Viewing distance was unconstrained, but averaged approximately 60 cm. Across both phases, a central fixation cross (0.70° x 0.70° of visual angle) and black ring (diameter: 19.93° of visual angle, thickness: 0.09° of visual angle) remained constant on the screen throughout each trial. Participants were instructed to maintain fixation during the trials. Even though fixations were not monitored during testing, all participants were trained to maintain fixation using methods developed by [3]. Each of the 7 target and non-target items subtended 1.74° of visual angle in diameter, and were encircled by white outlines that had a thickness of 0.09° of visual angle. Red square outlines (2.80° x 2.80° of visual angle, thickness: 0.26° of visual angle) were used to indicate which of these moving items were to be tracked during the trial. The colors of target and non-target items alike were sampled from three equiluminant colors (blue, pink, yellow), such that three items belonged to one color set, three items belonged to another color set, and a singleton was filled with the remaining color. Participants were informed that the colors of the items were uninformative during the thresholding phase, but that they aided the identification of each dot’s role within the motion structures during the test phase. After each trial was complete, participants received feedback in the form of red square outlines (2.80° x 2.80° of visual angle, thickness: 0.26° of visual angle) appearing around the correct targets.

#### Prediction task

In the prediction task, the location of two temporarily occluded target dots (green and red) had to be predicted at the end of the trial, using the motion of five (always visible) non-target dots. Stimuli and graphical user interface were written in custom Python code (using Matplotlib on the Qt5 backend) and were presented on an iMac16,2 (display resolution: 2048 × 1152; frame rate: 50 frames/second) in an anechoic experiment room at Harvard Medical School. Viewing distance was unconstrained, but averaged approximately 60 cm. In each trial, a gray circle (diameter: ∼12° of visual angle) remained constant on the screen throughout the trial. On the circle, seven dots (5 non-targets and 2 targets; diameter: 0.5° and 0.7° of visual angle respectively) moved stochastically according to eq. [4]. Dots were colored as indicated in **Fig. 10**: light-blue for dots 1, 2 and 3; light-green for dots 4 and 5; green for dot 6 (a target dot); and red for dot 7 (the other target dot). Participants were informed about the meaning of the colors within the motion structure of the respective block. At the trial end, participants predicted the targets’ locations by directing a green / red mouse cursor to the predicted location on the circle. The order (green / red) of the predictions was randomized across trials. After each trial, the true dot locations were revealed and participants received points based on the accuracy of their prediction:

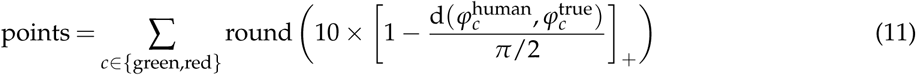

where d(·, ·) ∈ [0, *π*] denotes the (absolute) distance on the circle, and the rectification [·]_+_ restricts the points to positive values. This is a linear decay, per dot, from 10 points for a perfect prediction down to 0 points for errors of 90° and above. Note that due to the intrinsic stochasticity of the targets’ motion, even an ideal observer will usually obtain less than 20 points per trial. Participants received a bonus of $1 per 500 points (rounded up to the next dollar), resulting in $7 or $8 of paid bonus (in addition to the $10 per hour base pay).

### Discussion of the quantitative mismatch in the GLO condition of the MOT task

The computational observer model systematically underestimated human MOT performance in the GLO condition (cf. Fig. 2B in the main paper). In general, it is not surprising that a minimal observer model, that only contains the computational components essential to solving the MOT task, cannot capture all nuances of human tracking behavior. Nonetheless, we asked what is special about the global motion stimulus condition? First, we note that all motion conditions have the same marginal dot distance distribution because lim_*t →*∞_ *p*(***z***_*t*_) = ∏_*k*_ Uniform(*z*_*t,k*_; 0, 2*π*) for all conditions. However, dot proximity events, that is, times at which two dots have a small distance, typically last longer for GLO stimuli than for other stimulus structures due to the stronger positive velocity correlation of these stimuli. This is quantified in **Fig. 4** in terms of the auto-correlation function of pairwise dot (circular) distances. When two dots come close in the GLO condition, it takes several seconds until their locations are “decoupled” again. This differs in the other stimulus conditions.

Proximity events that span many video frames (each frame lasts for only 1/60Hz = 17ms) are expected to be particularly susceptible to *observation noise* in our observer model for two reasons. First, the employed DPVI algorithm [4] evaluates (and possibly discards) alternative mental dot assignments on a frame-by-frame basis and, thus, alternative hypotheses may not be maintained throughout a longer proximity event. Humans, in contrast, may reason on longer time scales. Second, the tracking process relies on Kalman filtering (forward inference) rather than smoothing (forward-and-backward inference). Updating past location estimates based on new observations could help to resolve ambiguous mental assignments in the recent past as they occur during proximity events. To test for a specific detrimental effect of observation noise to tracking of GLO stimuli, we re-evaluated the MOT data by decoupling the noise *σ*_obs_ assumed by the observer model and the noise *σ*_*x*_ actually corrupting observations ***x***_*t*_ in computer simulations (in Fig. 2B of the main paper, we had *σ*_obs_ = *σ*_*x*_ = 0.05). As an ad hoc-choice, we assumed that *σ*_obs_ = 10 *σ*_*x*_, such that corrupted observations in individual frames have a smaller weight in the Kalman update. Readjustment of the noise level to approximately reach the desired average target performance (2.15) in the IND condition yielded *σ*_obs_ = 0.7 and *σ*_*x*_ = 0.07. The resulting predictions of the observer model, using the decoupled noise for all stimulus conditions, are shown **Fig. 5** (additional analysis; only one trial repetition simulated). The resulting predictions match human behavior better than the computational observer model with *σ*_obs_ = *σ*_*x*_. Yet, even with decoupled observation noise, motion structured priors are required to explain human performance.

Surprisingly, stimuli with few, long lasting proximity events are potentially less susceptible to *decision noise* than stimuli with many short proximity events. In the context of our MOT model, decision noise refers to stochasticity in the mental dot assignments, as contrasted with observation noise that affects the perceived dot locations. Consider two hypothetical observer models which are characterized by having the same average performance in the IND condition: one model features a mix of small observation noise and decision noise, the other model features only observation noise, but no decision noise. During extended periods with two dots overlapping, a typical feature of the GLO condition, dot assignment might become highly ambiguous and unresolvable—independent of the presence/absence of decision noise. Outside of these periods, however, the smaller observation noise might benefit the first model. This computational argument could explain the relative performance gain of the Weber’s law observer, which features such mixed noise mechanisms, over the computational observer, which features only observation noise, in the GLO condition, independent of the employed prior (cf. Fig. 2B and 3B in the main paper).

We emphasize that the above discussion can only be a first step towards understanding the detailed contribution of observation noise and decision noise to multiple object tracking of motion structured stimuli. In addition to these algorithmic considerations, human participants may direct their visual focus and attention specifically to locations of proximity events in order to locally reduce observation noise [5]. However, teasing apart these different contributions was not the main focus of this study, which is why we did not pursue their study in further detail.

## Movie legends

**File:** `Movie_S1_MOT_IND.mp4`

**Legend: Example of the MOT experiment with independent motion**. This movie shows a screen recording of a trial in the independent (IND) motion condition.

**File:** `Movie_S2_MOT_CNT.mp4`

**Legend: Example of the MOT experiment with counter-rotating motion**. This movie shows a screen recording of a trial in the counter-rotating (CNT) motion condition.

**File:** `Movie_S3_prediction_GLO.mp4`

**Legend: Example of the prediction experiment with global motion**. This movie shows a screen recording of three trials in the global (GLO) motion condition.

---

**Figure 1.**
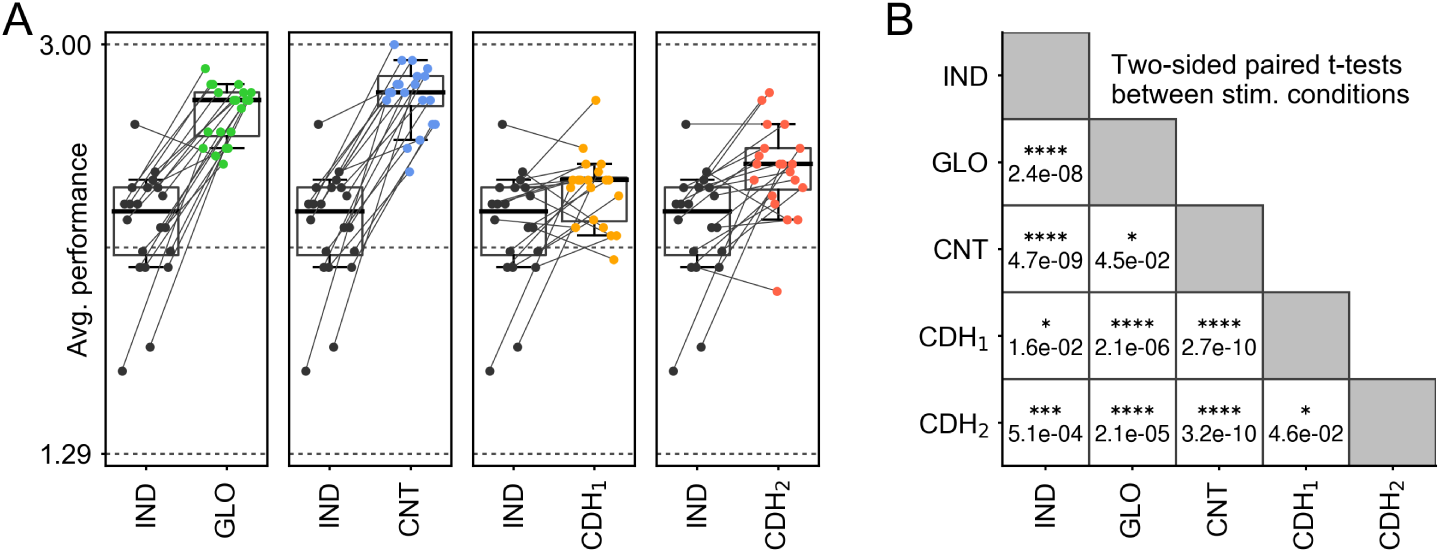
Pairwise comparison of human MOT performance. (A) Per participant performance in all conditions relative to the IND condition. The boxplots mark 10%, 25%, 50%, 75%, and 90% percentiles, respectively. The gray lines connect each participant’s performance across conditions. (B) Pairwise two-sided t-tests, testing for average performance similarity across all stimulus conditions (cf. Fig. 2B in the main paper, where one-sided tests were used for the specific hypothesis of performance increase).

**Figure 2.**
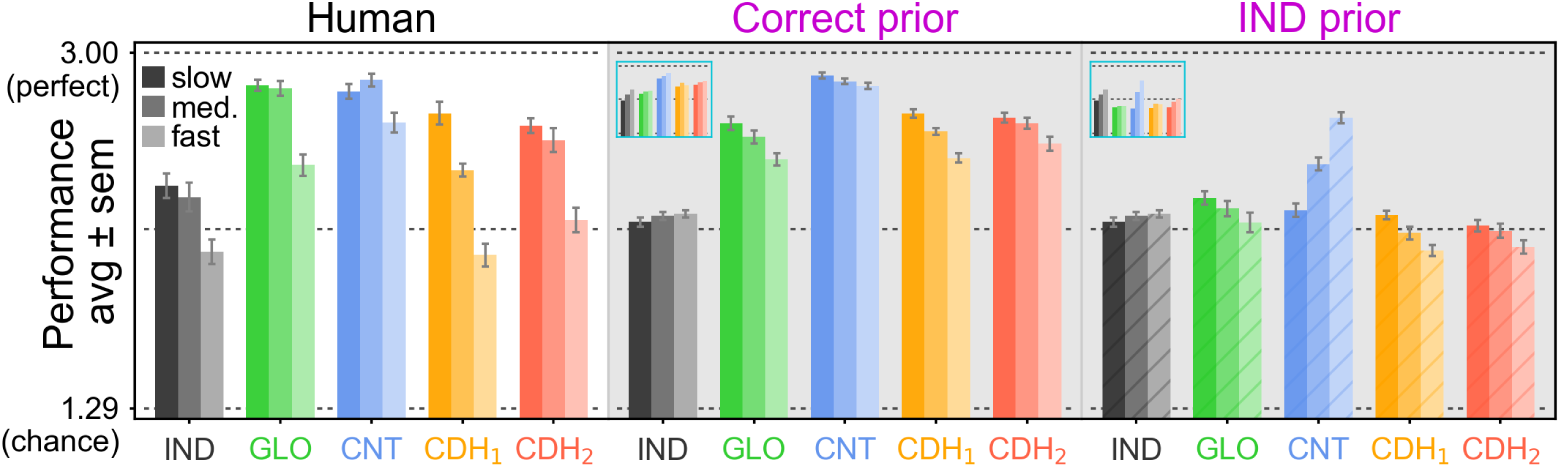
Speed-dependence of MOT performance. For each participant and condition, the 30 MOT trials were sorted by average dot speed (average over time and dots) and then assigned to three categories (slow, medium, fast; 10 trials each). Shown is the performance for each category, averaged over participants. *Left:* Human tracking performance declines at higher speed. *Center:* The Weber’s law observer features a qualitatively similar speed-dependence when the correct motion structure prior is employed, albeit not to the same amount. *Right:* If an IND prior is used, the speed-dependent decline and most of the general performance gain are lost. *Insets:* Same for the computational observer model, which does not feature velocity-dependent observaton noise.

**Figure 3.**
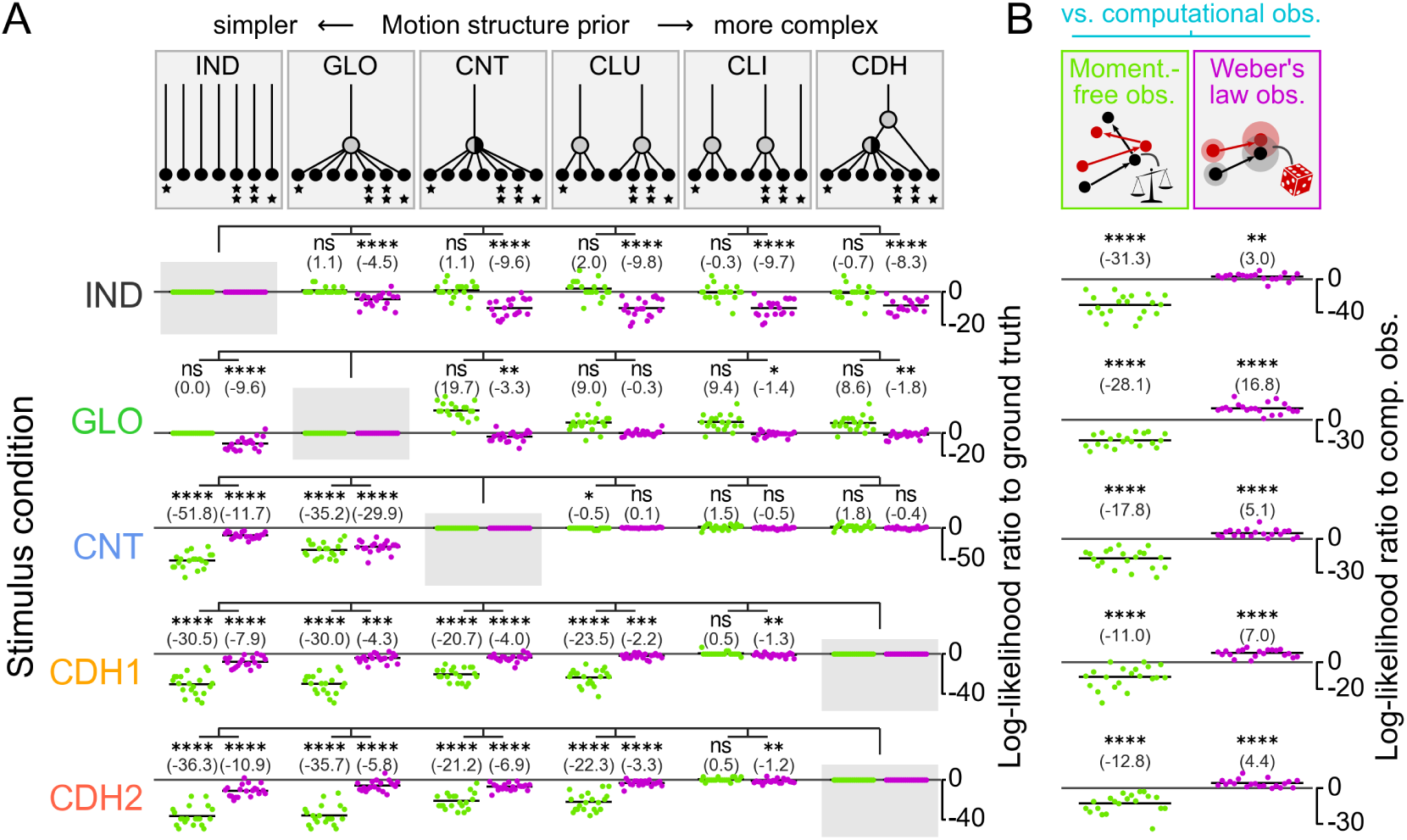
Bayesian comparison of employed motion structure priors in MOT for the alternative observer models. (A) Same as Fig. 3C in the main paper, but for the momentum-free (light green) and Weber’s law (magenta) observers. The results for the Weber’s law observer corroborate the finding that humans employ correctly or close-to-correctly structured priors during tracking. *‘s indicate significance of one-sided t-tests against the correct prior. (B) Comparison of the alternative models against the computational observer, when both models employ the correct motion structure prior. The Weber’s law observer explains human responses consistently better than the computational observer, especially in the GLO condition. The momentum-free observer, in contrast, cannot explain human MOT. *‘s indicate significance of two-sided t-tests against the computational observer.

**Figure 4.**
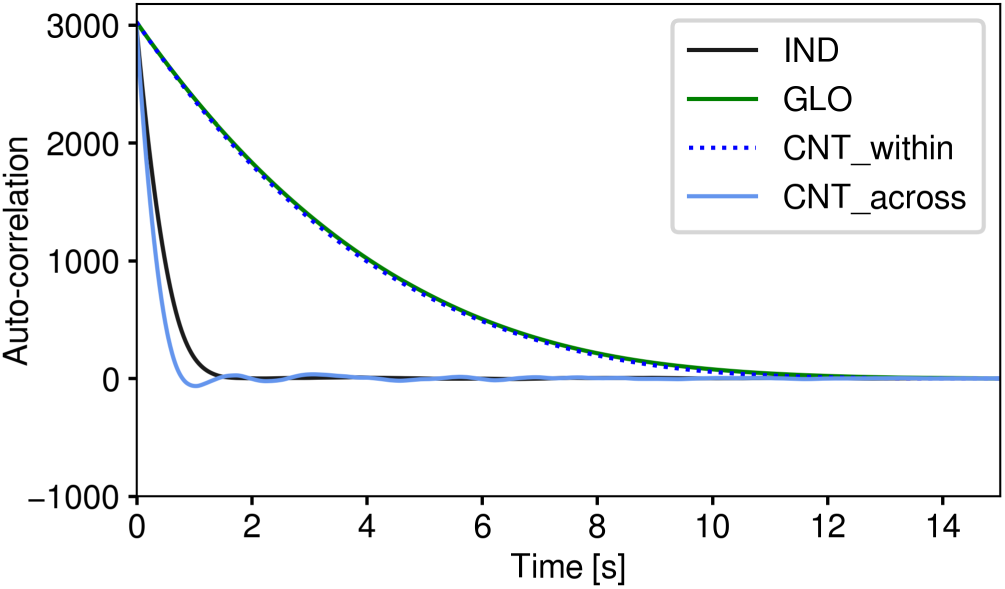
Auto-correlation function of pairwise dot distances in the MOT stimuli. All stimulus conditions share the same stationary location distribution *p*(***z***_*t*_) and, thus, also share the same distribution of dot distances. Here, we probe the temporal structure of dot distance in terms of the auto-correlation function. For the IND stimulus, dot distances become uncorrelated within a few hundred milliseconds. For the GLO condition, in contrast, dot distances remain mostly stable for multiple seconds. While GLO stimuli produce less events of “close encounter”, the events typically last longer, spanning many consecutive frames for making false dot assignments. A Markovian observer will not be able to resolve these errors later. For counter-rotating (CNT) stimuli, correlations are long only for pairs of dots within the same sub-group (CNT_within). Their confusion often does not impact performance for our target set. For dots from different sub-groups (CNT_across), in contrast, proximities last much shorter.

**Figure 5.**
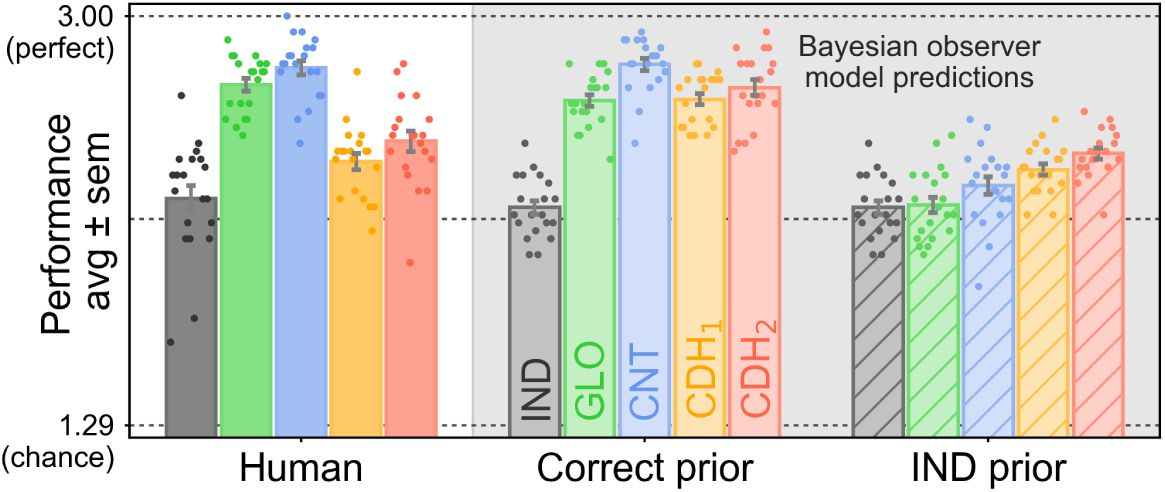
Probing a specific detrimental effect of observation noise to tracking of GLO stimuli. Re-evaluation of the MOT data with decoupled observation noise (for details, see *SI Appendix*, section “Discussion of the quantitative mismatch in the GLO condition of the MOT task”). Tracking of GLO stimuli is more affected by noisy observations than other stimulus conditions (especially, IND and CNT), likely due to the longer duration of dot proximity events.

**Figure 6.**
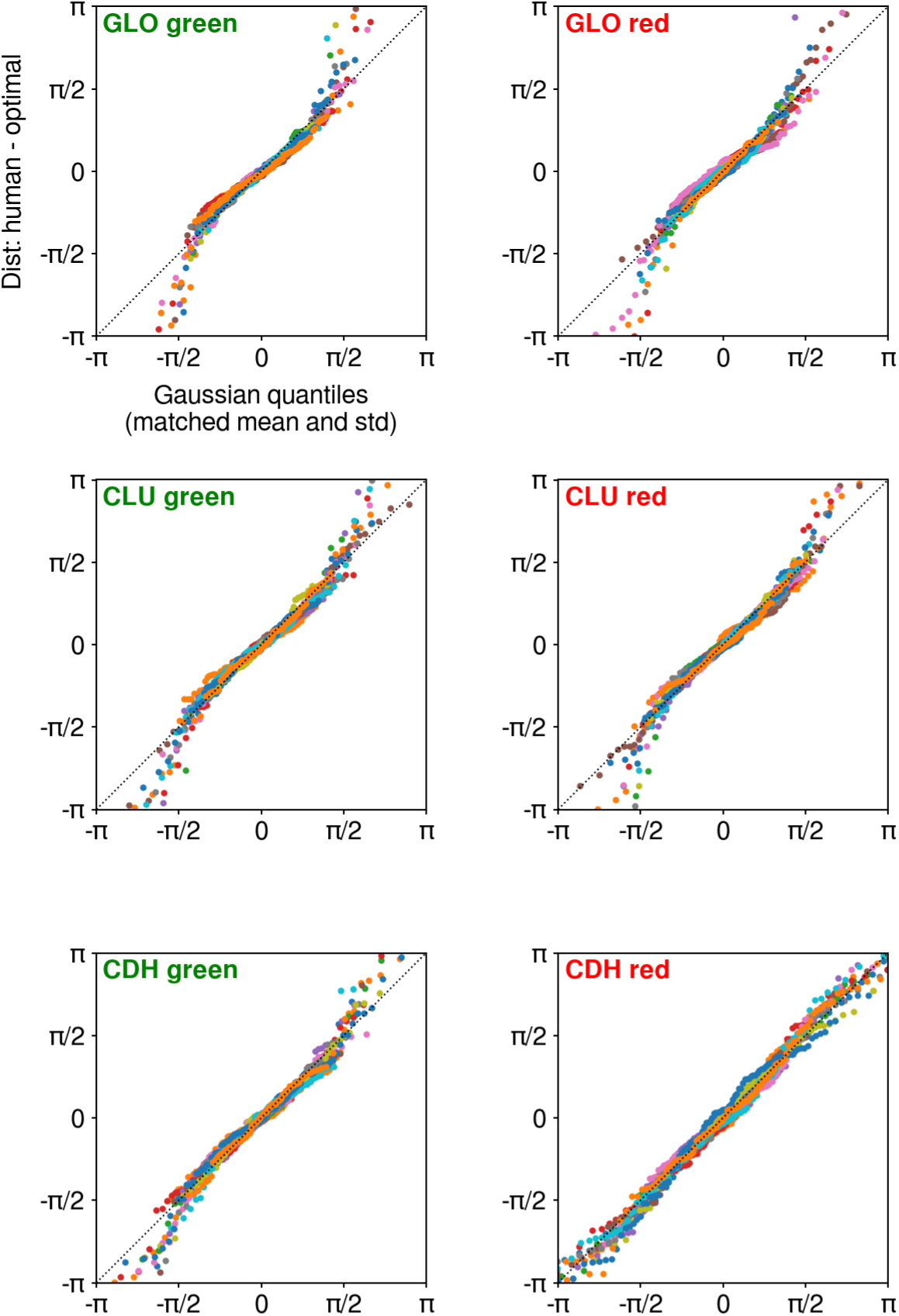
Human prediction suboptimality is normally distributed in the prediction task. Each panel shows a probability plot of prediction suboptimalities in a stimulus condition (GLO, CLU, CDH) for the green or red dot (one point per trial; colors=participants). Prediction suboptimality is the circular distance to the optimal prediction which is given by the mean ***µ***_kal_ of a Kalman filter with correct motion structure prior ***L***. Prediction suboptimalities (y-axis) are compared against a normal distribution with same mean and variance (x-axis; matched per participant). Most of the responses are along the identity (dotted line) as expected for a normal distribution. The small number of deviations at the tails can be attributed to dot confusion in some trials: swapping the red and green dot leads to uniformly distributed prediction errors, i.e., “heavy tails”.

**Figure 7.**
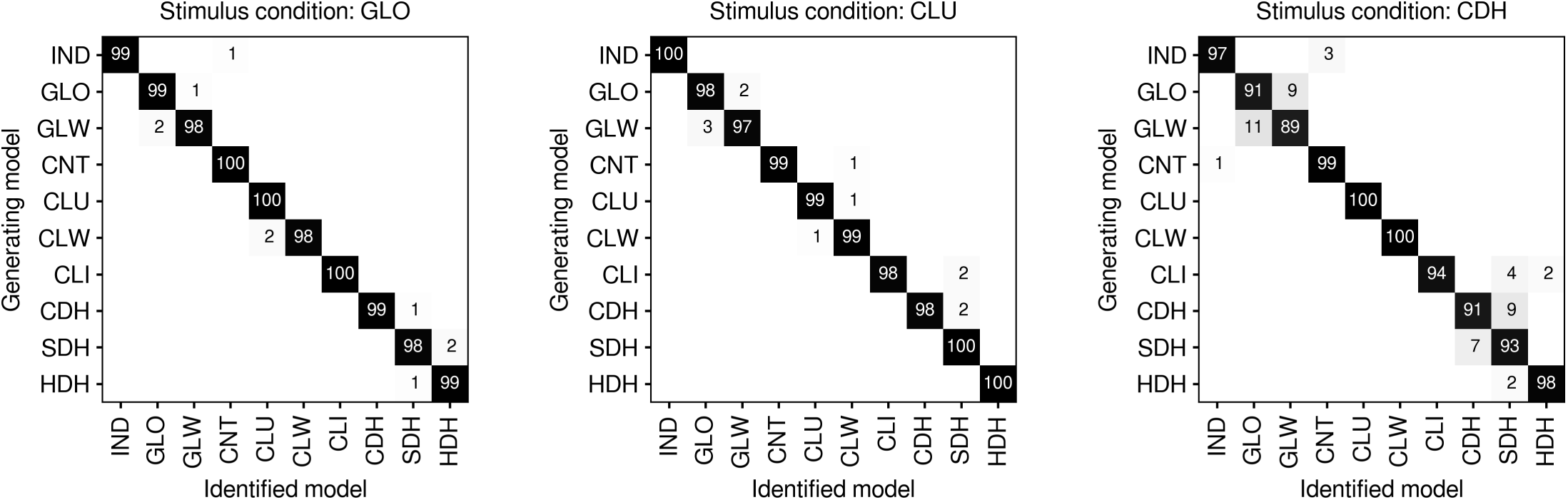
Fitting the Bayesian observer model can re-identify the generating motion prior in the prediction task. Per stimulus condition (GLO: left, CLU: center, CDH: right), we generated artificial responses to all 1200 (= 12×100) trials presented to human participants by drawing samples (*φ*_green_, *φ*_red_) from the observer model presented in *Material and Methods* for each of 10 candidate motion priors ***L***. The 10 priors are the 7 priors from the main paper and 3 additional priors (GLW: weak global, CLW: weak cluster, HDH: half deep hierarchy, see Fig. S10 for their definition). To create “human-like” responses, we used the average of the max. likelihood parameters *θ* = (*π*_s_, *a, b*) fitted to human participants under the correct prior ***L***^⋆^. Then we fitted all 10 candidate priors to the responses generated from the 10 priors as decribed in *Material and Methods*. This process (response generation and fitting) was repeated 100 times, i.e., in total 3×1200×10×100 = 3,600,000 artificial trials were evaluated. The figure shows the number of times, prior “Identified model” had the highest log-likelihood ratio in explaining the responses generated from prior “Generating model”.

**Figure 8.**
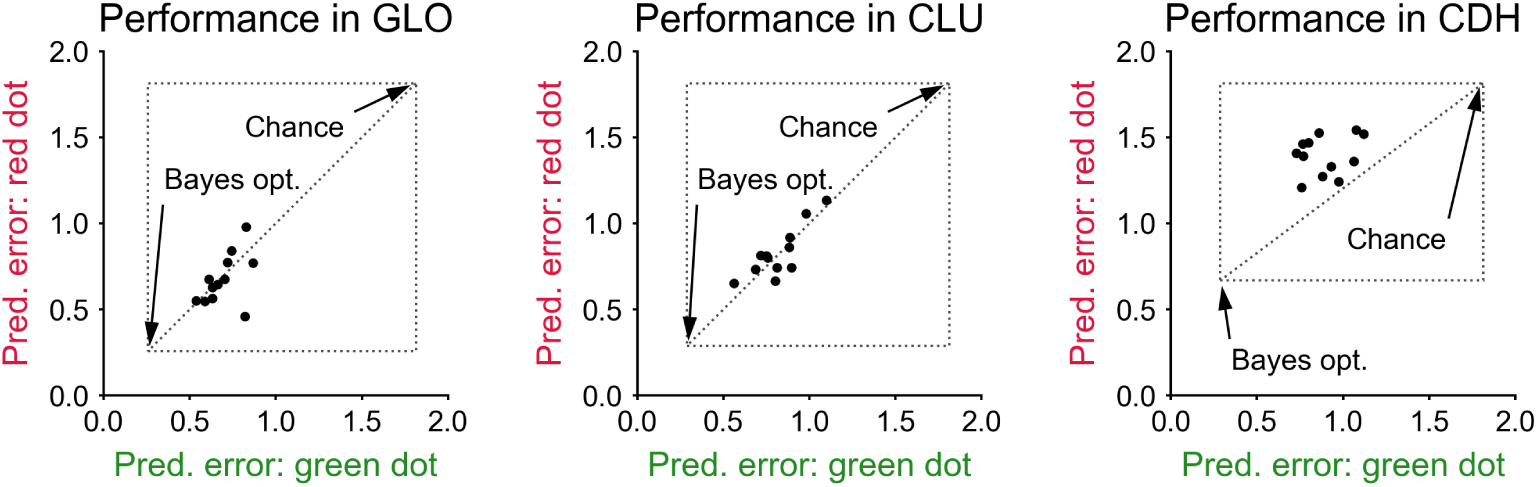
Human average performance in the prediction task. Same as Fig. 4B from the main paper, but for all motion conditions. In the CDH condition, the red dot’s location is significantly harder to predict, owing to its weaker coupling to the remaining dots.

**Figure 9.**
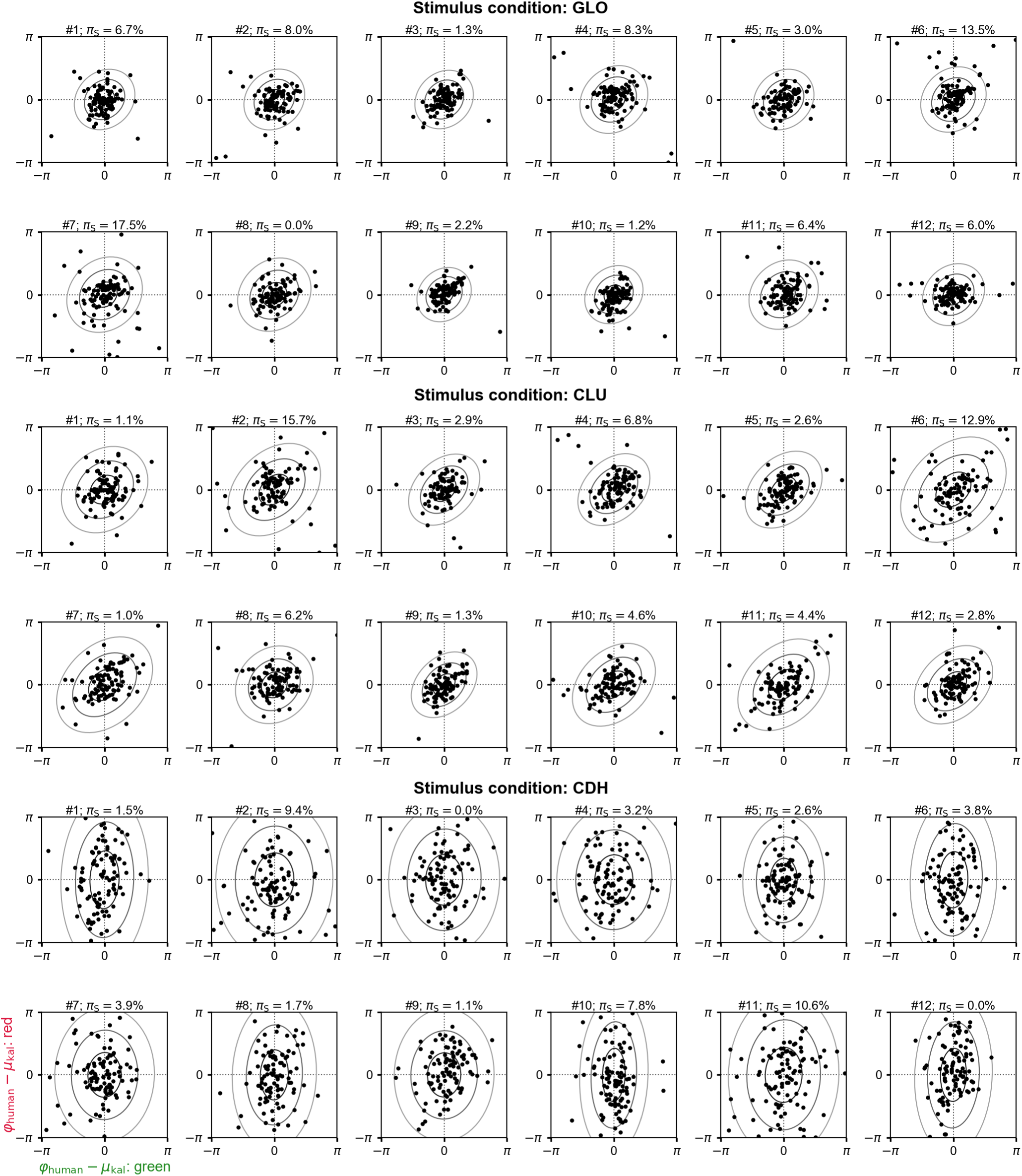
Model predictions of human responses in the prediction task. Same as Fig. 4C of the main paper, but for all participants and stimulus conditions. The fitted swap probability *π*_S_ is given in the panel titles. Note that only the non-swapped mixture component is plotted by the ellipses.

**Figure 10.**
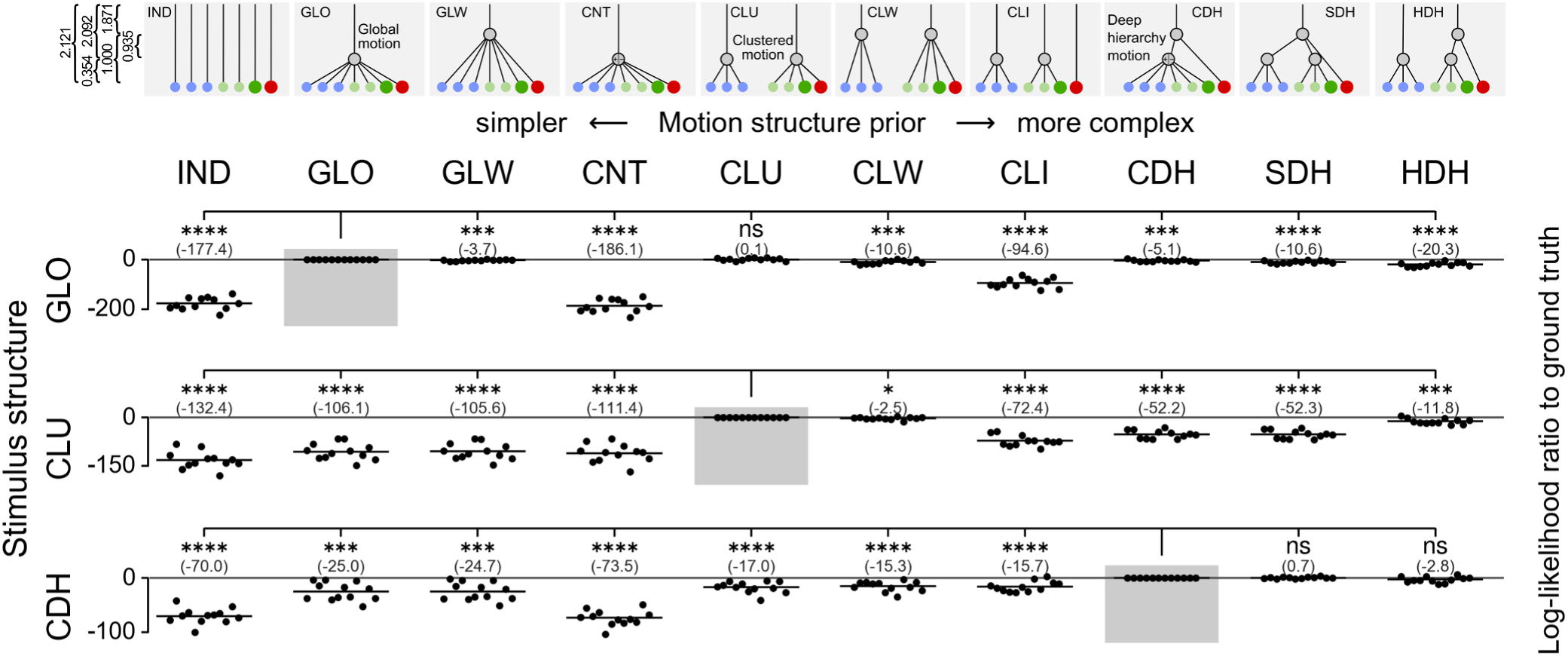
Revealing human motion priors in a multiple object prediction task (more candidate models). Same as Fig. 4E from the main paper, but with 3 additional (non-preregistered) motion structure priors, GLW: weak global, CLW: weak cluster, HDH: half deep hierarchy. These were included as additional, potential strategies to solve the task in the CDH condition. For instance, GLW mirrors the strength of the global motion component in CDH while disregarding the counter-rotating substructure. While GLW and CLW cannot explain human responses in the CDH condition, HDH – another nested structure with 3 latent sources – could explain human responses as well as the correct CDH prior. Due to the similarity of GLO and GLW (or CLU and CLW), these priors could not be distinguished reliably in the GLO (CLU) condition. Motion strengths *λ* (vertical extend of edges in the graphs) are not drawn to scale. Numerical values of all motion strengths *λ* are given in the top-left corner of the figure. For our stimulus design, motion strengths add quadratically, granting control over marginal dot velocities. At the example of GLO motion, 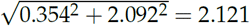 matches the strength of IND.

**Figure 11.**
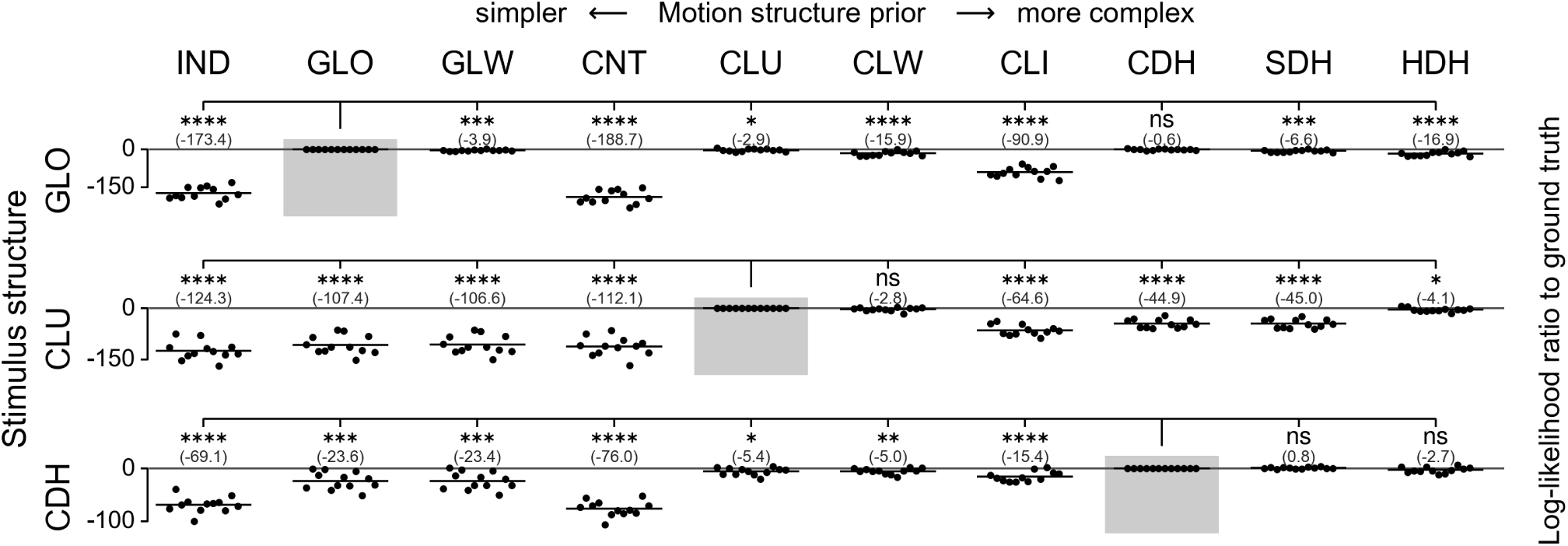
Benefits of multiple object prediction over single object prediction. Same as Fig. 4E from the main paper (and Fig. S10), but with an observer model that disregards the posterior covariance **Σ**_kal_ by modeling human responses as

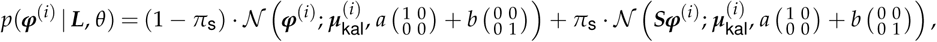

i.e., replacing the structured covariance by unstructured noise for both dots (note that the model still has 3 free parameters). This evaluation resembles the type of data that could be measured in a single object prediction protocol where the green and red dots’ locations are predicted in separate trials. Compared to our experiment, it would thus require twice the number of trials. Despite this, a general reduction of contrast in the log-likelihood ratio is observed, when compared to multiple object prediction (Fig. S10). This reduction is especially apparent in the CDH stimulus condition where differentiation of the CLU tracker is reduced to the edge of statistical significance.

**Figure 12.**
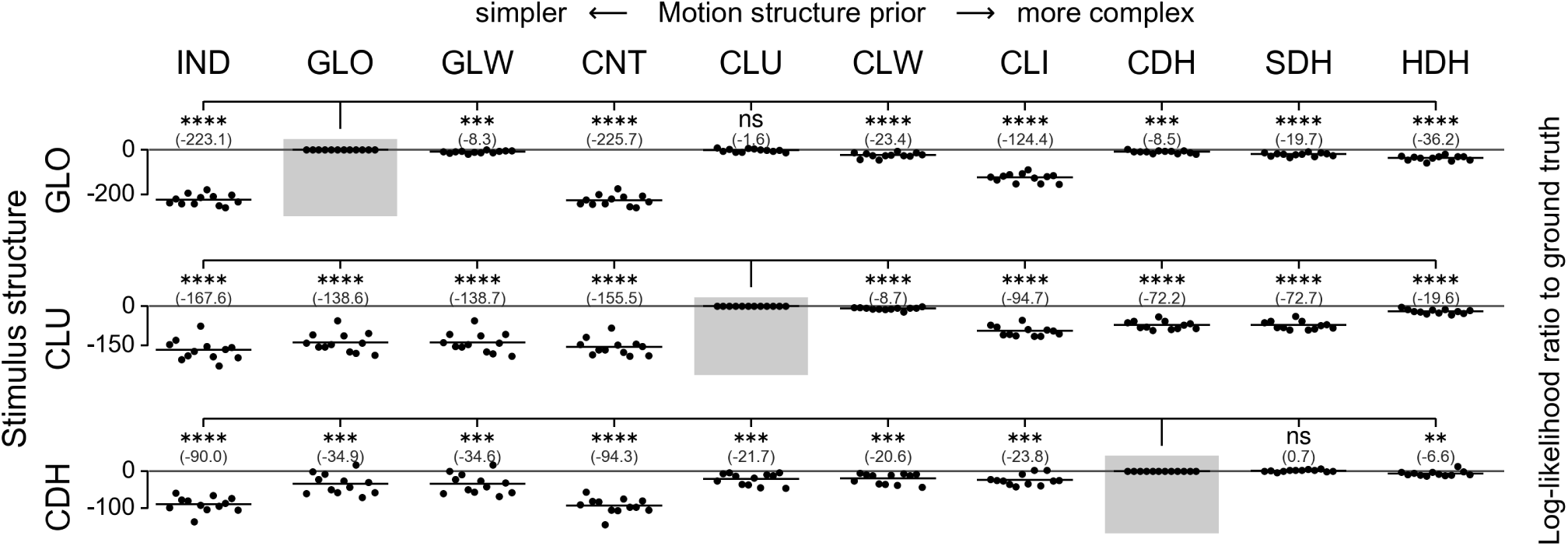
Explaining the prediction task with a momentum-free observer. Same as Fig. 4E from the main paper (and Fig. S10), but using the dynamics of the momentum-free observer model (w/o dot confusion) for making predictions. Solving the prediction task critically relies on exploiting motion relations between the dots, but does not necessitate the concept of inertia.

**Figure 13.**
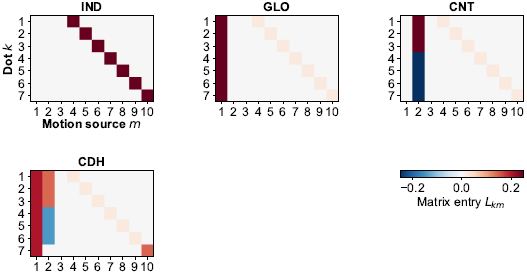
Motion structure matrices of the MOT task. All matrices ***L*** lead to the same marginal velocity distributions *p*(*v*_*k*_) for all dots *k*. In this plot, we set *f*_speed_ = 2.0 (a typical value for human participants).

**Figure 14.**
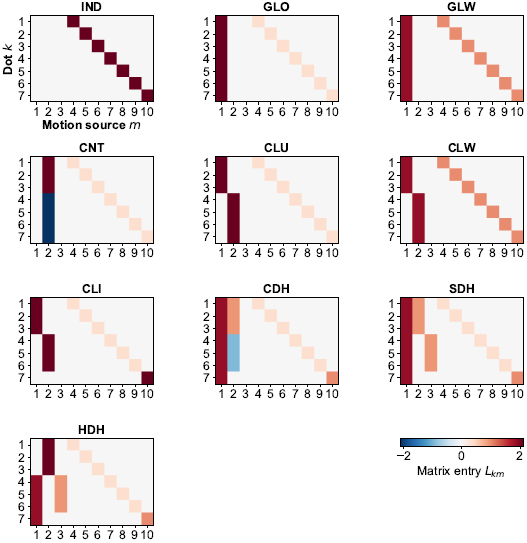
Motion structure matrices of the prediction task. All matrices ***L*** lead to the same marginal velocity distributions *p*(*v*_*k*_) for all dots *k*.

